# Robust Multi-Mutant Protein Stability Prediction from a Fine-Tuned Evolutionary Scale Model

**DOI:** 10.64898/2026.06.04.730231

**Authors:** Shawn Reeves, Subha Kalyaanamoorthy

## Abstract

Recently, high-throughput experimental techniques have propelled improvements in deep learning-based prediction of mutation effects on protein stability. However, leading stability predictors still struggle to predict the combined effect of multiple mutations and prefer mutations that negatively impact other properties, including expressibility. To mitigate these limitations, we apply Low-Rank Adaptation (LoRA) to specialize ESM3 for stability prediction by fine-tuning on the Megascale protease susceptibility dataset, developing a novel dual-perspective inference mechanism to provide explicit mutant context information. ESM-Mutant Stability Ranker (ESM-MSR) significantly exceeds all contemporary methods tested on the prioritization of stabilizing mutations (ΔNDCG@96 ≥ +0.12), double mutant ranking (Δ*ρ*_*avg*_ ≥+0.068) and direct epistasis ranking (Δ*ρ*_*avg*_ ≥ +0.164) within the Megascale test set. Further, it generalizes effectively to heterogeneous thermostability benchmarks, consistently matching or exceeding current approaches across our comprehensive suite. Finally, a single parameter *σ* enables tunable control of the model’s compromise between stability and more general sequence fitness, leading to state-of-the-art performance in the Human Domainome 1 benchmark (*ρ*_*avg*_ = 0.573) at *σ* = 0.5, demonstrating the broad applicability of ESM-MSR as a protein engineering tool.

## Main

Thermodynamic stability is a fundamental property of most proteins, as it is usually a prerequisite for maintaining a biologically active folded configuration. However, evolutionary pressure optimizes only the fitness of an organism, which is unlikely to improve through stabilization of any protein that can already remain folded under native environmental conditions [44]. Hence, natural proteins tend to be only marginally stable and, in order to retain function in non-native environments, stability-increasing modifications are often necessary. Introduction of stabilizing mutations to enzymes can enable high-temperature industrial catalysis [38] and rescue heterologous expression of misfolding-prone proteins [16]. Stabilized mutants can also serve as starting scaffolds with improved mutational tolerance when attempting to increase activity or substrate promiscuity (e.g. by directed evolution), since functional gains often entail a stability cost [43]. Finally, accurate prediction of mutant effects on stability can aid in understanding the nature of genetic diseases and protein evolution [23].

Despite its importance, the field of mutant stability prediction has historically been fraught with model bias and poor generalization, largely consequences of the limited diversity of experimental data and flawed testing methodology [11, 4]. The most robust among hundreds of published stability predictors tend to combine biochemical or physical knowledge with regression on experimental data. For example, Rosetta Cartesian DDG [15, 1] approximates the physical interactions of residues within a protein using interatomic potentials, but the parameters of the force-field are adjusted to better reproduce experimental stability measurements. Advances in self-supervised protein modeling have circumvented previous limitations by capitalizing on a wealth of diverse training data that can indirectly inform a model about stability. New pre-trained models demonstrate zero-shot (lacking task-specific training) mutant stability predictions that match or exceed the best bespoke stability models of the last generation, including Rosetta [24, 37]. Zero-shot predictions are rendered by comparing the modeled log-likelihood of the mutated residue identities to the wild-type identities; if the mutant identity is more likely, it may be more compatible with its protein context and hence more stable.

A promising emerging paradigm to improve predictive performance involves specializing these pre-trained self-supervised models via supervised transfer learning using experimental stability data. ThermoMPNN [9] exploited prior knowledge about the structure-conditioned likelihood of amino acid identities learned by ProteinMPNN [6] by training a stability prediction module to transform latent embeddings from that pre-trained model into stability predictions. To achieve this, they capitalized on the work of Tsuboyama et al., who dramatically expanded the available mutant stability data from a few thousand mutants sparsely sampled from hundreds of proteins to roughly 776,000 densely sampled mutants of small protein domains using their high-throughput protease susceptibility assay [45]. Their resulting “Megascale” dataset provided enough data for ThermoMPNN and other new models such as the AlphaFold2-based Mutate Everything [31] and the ESM2-ProteinMPNN hybrid SPURS [22] to learn to transfer the representations of their respective base models into robust mutant stability predictions.

Few stability predictors can model interactions between mutations due to a combination of computational limitations, training data scarcity, and the need for specialized architectures. However, these interactions can be crucial considerations in designing stable proteins. For instance, mutating a buried hydrophobic residue to a charged residue would generally be highly destabilizing, but a simultaneous mutation introducing a nearby opposite charge may partially offset this effect through the formation of a salt bridge. While all mutant stability models are theoretically capable of approximating multi-mutant effects by treating the mutations independently and simply adding their predicted effects, these non-additive “*epistatic*” effects occurring due to interactions between mutations are neglected. While Rosetta, alongside closed-source methods like Dynamut2, DDMut3D and DDGemb can directly predict multi-mutant effects, no evidence has yet been presented to suggest that these methods learn epistasis or outperform their own additive baselines [39, 49, 41], and prediction of the mutated structure (except in the latter) introduces a huge computational throughput bottleneck. An efficient deep learning approach involves encoding only the wild-type protein using a pre-trained model and then deriving and operating upon mutant representations in the latent space, but Mutate Everything and ThermoMPNN-D used this approach and failed to consistently improve upon their respective additive approximations [31, 8]. Since direct design of mutants with explicit consideration of epistasis could greatly reduce the experimental burden in developing stabilized proteins and expand the space of candidate mutants from hundreds to millions, this is a key challenge to overcome [3].

We sought to create an efficient epistasis-aware model by fine-tuning a cutting-edge multi-modal model on the Tsuboyama Megascale dataset. ESM3 integrates sequence, structure and function during pre-training and exhibits a strong understanding of the complex interactions between modalities through excellent zero-shot stability and function prediction capabilities [17]. Furthermore, at the time of writing, ESM3 is recognized as the third highest performing among nearly 100 distinct approaches for zero-shot stability prediction on the ProteinGym benchmark [29]. To leverage ESM3’s prior knowledge for stability prediction, we implemented supervised fine-tuning via Low-Rank Adaptation (LoRA) [20]. LoRA is a parameter-efficient approach that adapts a pre-trained model to a new task without modifying the original weights, based on the hypothesis that weight updates for adaptation have a low intrinsic rank. By introducing trainable low-rank decomposition matrices into the frozen layers, LoRA efficiently adapts the model using limited training examples and mitigates overfit. The parameter-efficient fine-tuning strategy has previously shown promise in various downstream prediction tasks in protein science [42]. In this work, we adapt the 1.4 billion parameter ESM3-small-open model using two distinct LoRAs totaling less than 1.3% additional parameters. The two LoRAs correspond to viewing a mutation from both the wild-type and mutant “perspectives”, where epistatic interactions are explicitly encoded in the latter. Masked language models including ESM3 are trained to predict the identity of hidden (masked) amino acids within a mostly unmasked sequence, and so all mutated positions are generally masked during zero-shot inference and the scores at each position are summed, known as the “masked marginal” scoring strategy [24]. However, since this approach does not provide explicit information about the interacting residue identities, we chose to adapt the model to use fully unmasked sequences. Because we achieve state-of-the-art mutant ranking performance on a wide collection of held-out test sets without compromising performance on related properties, we anticipate that ESM-MSR will be an effective tool for protein engineers and bioinformaticians alike. To facilitate its widespread usage in these fields, we open-source the code and provide a simple-to-use mutation scoring and visualization pipeline (see Methods).

## Results

### Overview of Approach and Test-Set Ranking Performance

We first showcase the ability of ESM-MSR to rank mutants of a given protein from most-to least-stabilizing after it is trained according to the procedure indicated in Figure 1a and elaborated in the Methods. We compare transfer learning methods, trained using the same data split, with Rosetta Cartesian DDG, ProteinMPNN, and ESM3 pre-trained models as static baselines. All methods are compared on an identical test set of 43,700 (32,968 single- and 10,732 double-) mutants of small protein domains with low sequence identity to any training or validation domain from the Tsuboyama Megascale dataset (see Supplementary Figure 1 for a visualization). Figure 1c&d demonstrate that ESM-MSR significantly (*p <* 0.002) outperforms all other transfer learning methods at (c) prioritizing and (d) ranking mutants with the same distribution as the training data. Fine-tuning is a key element to this performance, contributing an average 0.309 increase in NDCG@96 and a 0.256 increase in Spearman’s correlation over the zero-shot prediction from the pre-trained base model (ESM3-small-open; Overall), and newly enabling modest relative ranking of stabilizing mutants. The inability of pre-trained models to correctly rank stabilizing mutations is consistent with the absence of a fitness signal; increasingly stabilized proteins are seldom more fit, and thus their modeled likelihood is not higher than wild-type [44]. While the other transfer learning methods (SPURS, ThermoMPNN and Mutate Everything) also learn to rank and prioritize stabilizing mutations, they achieve only a fraction of the ability of ESM-MSR. In addition to traditional regression error minimization, ESM-MSR also uses a second fundamentally different training objective from the other models: per-protein rank-based optimization. Specifically, the ListMLE loss [47] strongly encourages ESM-MSR to identify stabilizing patterns, where regression losses must fit the whole distribution, mostly determined by moderately destabilizing effects. Although we suspected that this ranking loss was a key contributor to stability recovery, a mean squared error-only ablation did not show degradation in ranking stabilizing mutations (Supplementary Tables 1-3), suggesting that the effect instead originates from the architecture of ESM-MSR. Meanwhile, despite training directly on stability measurements, Rosetta’s overall performance lies somewhere between the stability-naive pretrained models and the supervised transfer learning approaches. Since self-supervised pre-trained models such as ESM3 are expected to preserve the functional characteristics of proteins better than stability models [37, 19], and Rosetta requires orders of magnitude more time to predict each mutant than any other tested method, the role of Rosetta in stability engineering shifts toward specific use cases such as interpretable modeling of side-chain interactions. Relative to the zero-shot pre-trained models, the four transfer learning methods built from self-supervised models (SPURS, ThermoMPNN, Mutate Everything, and ESM-MSR) are also expected to sacrifice some of their base models’ understanding of general protein fitness in order to improve stability prediction. We show evidence of this in later sections.

**Figure 1.**
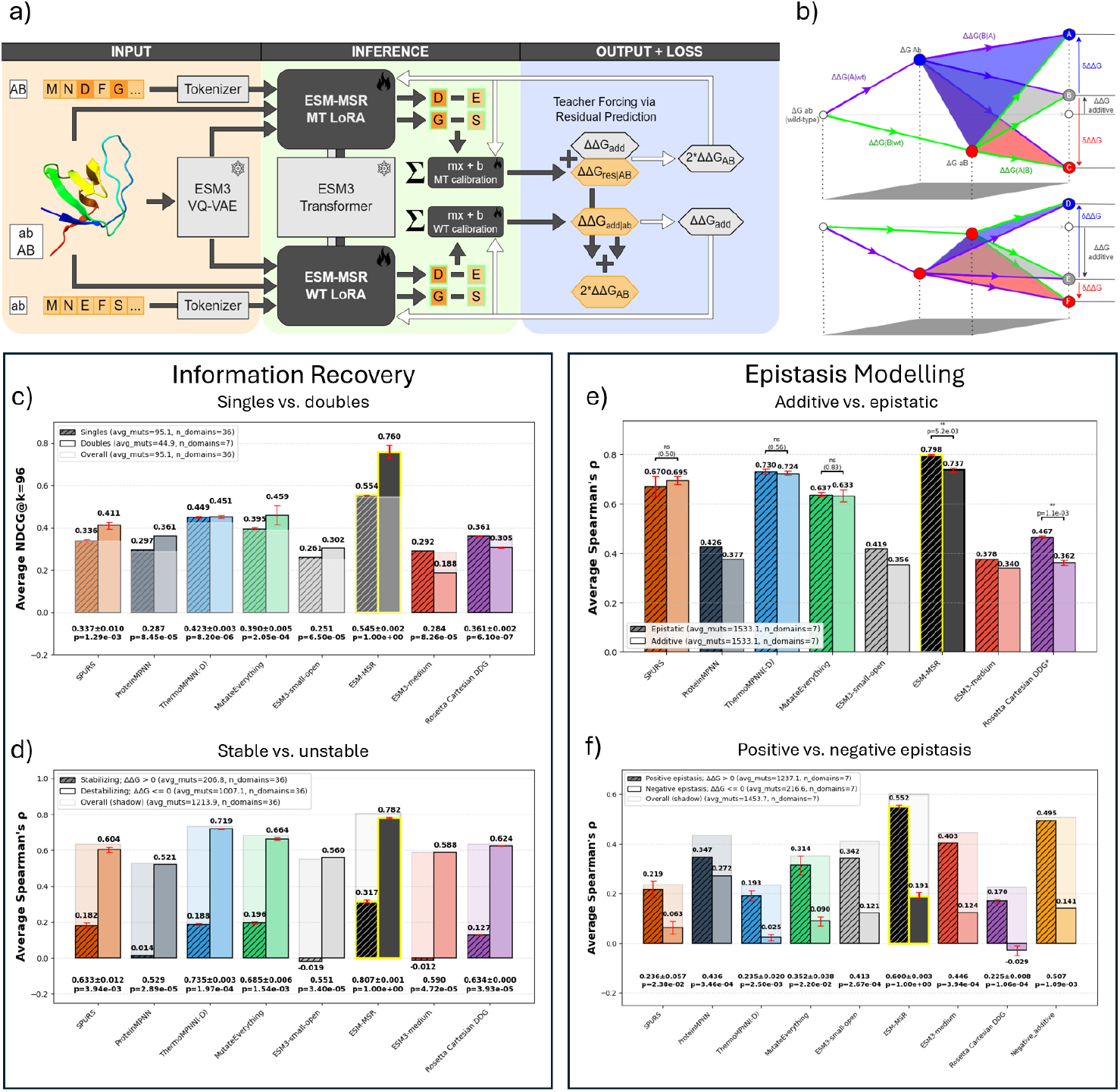
Overview of ESM-MSR Training and Performance on the Megascale Test Set. a): A simplified diagram of the training and inference processes for ESM-MSR. The mutated sequence is passed to the MT LoRA while the wild-type sequence is passed to the WT LoRA. Wild-type backbone coordinates and structure tokens generated using the VQ-VAE are passed to both branches. Both trainable LoRAs use the same frozen ESM3 backbone, but the LoRAs are independently applied and each shifts latent representations to be more stability-focused. ListMLE and mean squared error losses are used to update the weights of the LoRA and calibration head (white arrows); the WT LoRA is trained to predict the additive effect and the MT LoRA is trained by teacher forcing to predict the residual from additivity. b): Visualization of various types of epistasis in double mutants, inspired by Miton et al. [26]. ΔΔ*G* is defined relative to the wild-type unfolding free energy change, while *δ* ΔΔ*G* compares an observed effect to ΔΔ*G*_*additive*_. Double mutation can lead to additive effects between mutations (B,E), synergy (A,D), or clashes (C,F). c): Test set information recovery performance for the full set of 43,700 test set mutations, overall (background shadow / foreground ghost) and after splitting into single (hatched bars) and double (solid bars) mutant subsets. Average *NDCG*@96 uses the top 96 highest scoring mutants per model and assigns gains for stabilizing mutants only (otherwise, 0). Some domains have fewer than 96 true stabilizing mutants, tracked in avg_muts in the legend. For transferred models, error bars represent the standard deviation across three differently seeded training runs using the same data. For Rosetta, they represent the standard deviation in score across three pairs of mutant and wild-type structure models. *p* values represent the probability of the null hypothesis that the overall performance is the same as ESM-MSR; models with error bars are compared to ESM-MSR using a two-sample Welch’s t-test, while pre-trained models have only one sample and use a one-sided t-test. d)-f): Test set ranking performance (average Spearman’s *ρ*), using analogous visual elements. d) All test mutations, overall or split into stabilizing (ΔΔ*G >* 0) or destabilizing (ΔΔ*G* ≤ 0) subsets. e) Double mutants only, split into epistatic (native double mutant predictions) or additive approximation (sum of constituent mutation predictions). f) For 10,176 double mutants with both singles characterized, epistasis is derived from experimental or predicted ΔΔ*G*_*AB*_ − ΔΔ*G*_*A*_ − ΔΔ*G*_*B*_, overall and split into positive (*δ* ΔΔ*G* > 0) or negative (*δ* ΔΔ*G* ≤ 0) epistasis subsets.

### Double Mutants and Epistasis

We next evaluate the ability of ESM-MSR to capture non-additive (epistatic) effects of mutations. As shown in Figure 1b, double mutation can lead to various outcomes relative to the naive additive assumption of independent mutation effects, especially by synergizing or clashing with each other. Figure 1e shows that the “epistatic” double mutant predictions of SPURS, ThermoMPNN and Mutate Everything are not significantly more correlated to the ground truth than their corresponding additive approximations, indicating that each has failed to correctly model non-additive effects. Possible mechanisms for this shortcoming are examined later in this section and have important implications for training future transfer learning stability models. In contrast to the two other transfer learning methods, ESM-MSR gains a statistically significant (Welch’s t-test, *p* = 0.0052) and sizable (Δ*ρ* = 0.061) advantage by considering epistasis, comparable to the performance delta of its base model between prediction strategies. The benefit conferred by modeling interacting mutations is greatest for Rosetta, which benefits from explicit physical modeling and backbone conformational sampling [33]. Specifically, the structural relaxation performed by Rosetta allows Cartesian DDG to assess a more plausible, mutant-accommodating structure while all remaining methods use only the wild-type coordinates. ESM3 is also augmented by considering non-additive interactions between mutations using our dual-perspective inference scheme, but applying our dual-view approach to ESM3 does not outperform the traditional masked marginal scoring approach. It appears that fine-tuning is required to enable the model to escape its masked language modeling pre-training objective and learn to score unmasked positions, and ESM-MSR outperforms ablations trained to use masking (Supplementary Tables 1-3).

To directly probe the model’s understanding of non-additivity and interaction, we also assessed predicted epistasis (*δ*ΔΔ*G*, the non-additive interaction between mutations defined as ΔΔ*G*_*AB*_ − (ΔΔ*G*_*A*_ + ΔΔ*G*_*B*_)) in Figure 1f. Zero-shot predictors evidently have a strong prior understanding of stability epistasis, with ESM3 and ProteinMPNN overall outperforming all methods except ESM-MSR. Interestingly, supervised approaches (including Rosetta) are essentially incapable of ranking the negative epistasis subset, meaning that these models cannot distinguish relative incompatibilities, such as steric clashes and repulsion. This is especially surprising for ThermoMPNN and SPURS, which use representations derived from the strongest model for this task, ProteinMPNN. For transfer learning models, the effect is related to survivorship bias in the training and testing data: mutants with large destabilizing effects fall outside the dynamic range of the experiment and are excluded. Remaining double mutants, statistically arising from mostly destabilizing single mutants, must exhibit compensatory positive epistasis to “survive” and have effects in the measurement range. Consequently, negative epistasis is artificially rare (leading to the imbalanced categories seen in the legend) and positive epistasis is strongly anti-correlated with additive stability, shown with the Negative_additive baseline. Furthermore, double mutants with additive effects below the assay floor (≈ −3.5 kcal/mol) have epistasis effects that are strongly anti-correlated with additive stabilization and are very hard to rank accurately for models not trained on this dataset (Supplementary Figures 5&6). Models trained on the Megascale data can overfit by learning that increasingly destabilizing single mutants should make less destabilizing contributions in double mutants, exhibiting compensatory epistasis (the range between D and E in Figure 1b). MutateEverything appears to fall victim to this bias more than the other transfer learning approaches (Pearson’s *r* = −0.6 between its epistasis predictions and ground truth additive effects, Figure 2a), probably because it lacks the over-and-back augmentation used by ThermoMPNN-D (*r* = −0.09) and the strong statistical priors of SPURS (*r* = −0.19) and ESM-MSR (*r* = 0.00). We also performed density capping (see Methods) to mitigate this behavior. Survivorship bias may influence pre-training as well: zero-shot predictors perform much worse in ranking negative epistasis, perhaps because few examples of conflicting residues appear in sequence databases. This failure suggests that all studied models (including ProteinMPNN, whose performance degrades on an alternative test set in Supplementary Figure 7) are unable to anticipate the structural consequences of interacting mutations. On the other hand, ESM-MSR’s exceptionally strong positive epistasis ranking may reflect pairs of mutations that are well-accommodated into the wild-type backbone configuration, leading to the very high NDCG seen in Figure 1c and positioning it as an excellent tool for double-mutant engineering. Viewing the problem instead as a classification between positive and negative epistasis (Supplementary Figure 8) shows that all models struggle to differentiate the two classes, with ESM-MSR performing best (MCC = 0.128) followed by Rosetta (MCC = 0.109).

**Figure 2.**
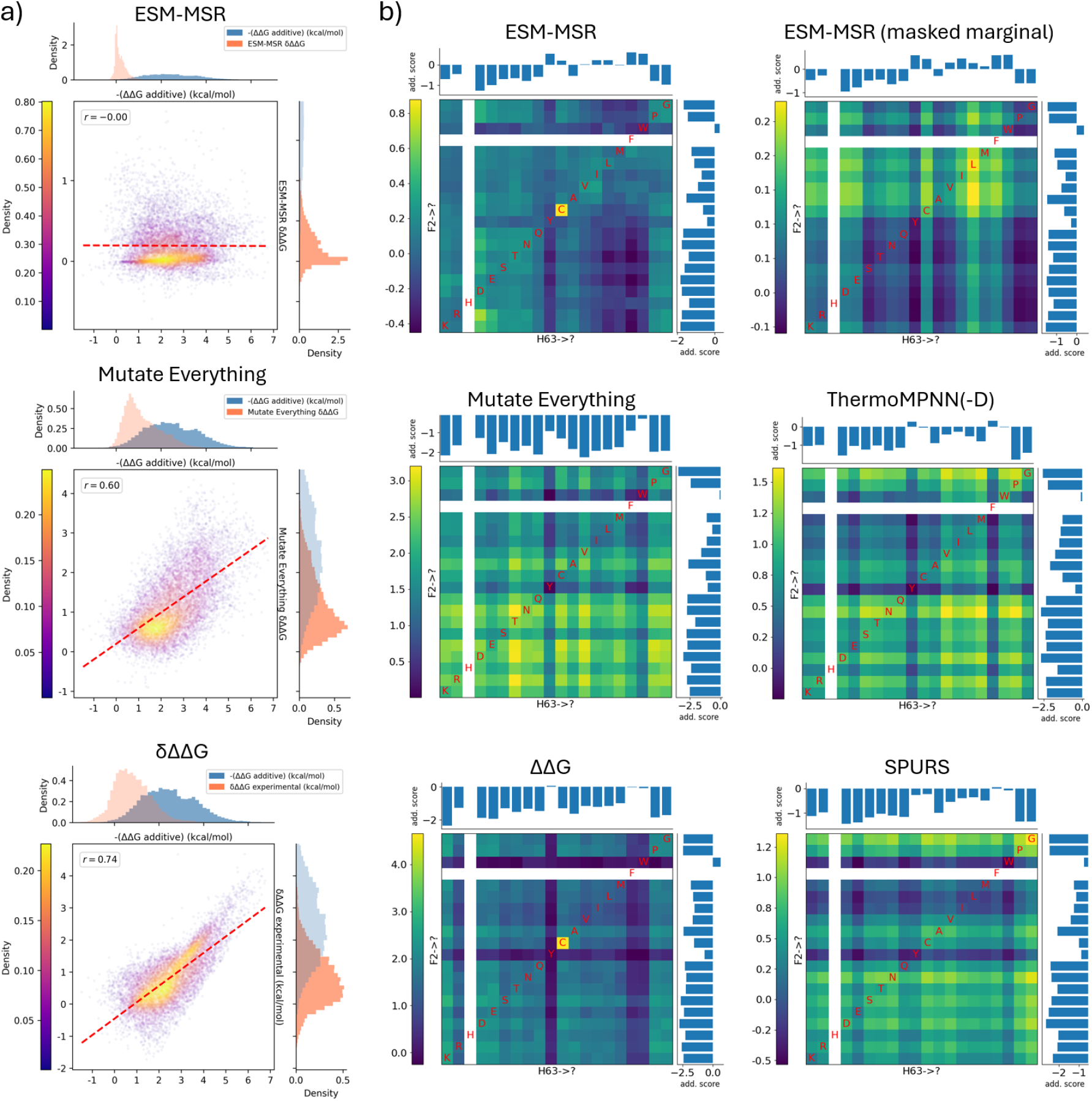
Inspection of Epistasis Modeling Behaviors. a) Correlations between the additive ΔΔ*G*_*A*+*B*_ label and the predicted epistasis for ESM-MSR, MutateEverything, and the ground truth *δ*ΔΔ*G* label. Point color indicates density (kernel density estimate) and the dashed red line indicates the linear best fit. The Pearson correlation is given in the top-left annotation. Marginal plots indicate the distributions, with the darker distribution indicating that axis’ plotted distribution. b) Heatmaps showing the pairwise epistasis for every characterized pair. White rows and columns are the wild-type identities. Colors indicate predicted epistasis, with yellows indicating more positive epistasis and blues indicating more negative epistasis. One-letter amino acid identities are in red on the diagonal. Marginal distributions show the scores of the corresponding independent single mutants.

SPURS and ThermoMPNN-D’s lower epistasis ranking performance may also be explained by their failure to model *identity-dependent* pairwise epistasis, instead learning only approximately *identity-independent* epistasis. In this context, we define identity-dependent epistasis as the component of total epistasis that depends on the compatibility of a specific pair of amino acids, contrasting the “naive” component that considers only how much more or less stabilizing a substitution should be in the context of *any* partner mutation. The nature of this effect is made clear in Figure 2b, where SPURS’ and ThermoMPNN-D’s (and to a lesser degree, Mutate Everything’s) behavior manifest as predicted pairwise interaction matrices with “plaid” patterns; relative to single-mutant predictions, both models merely apply a nearly fixed offset to each constituent single mutation of a double. Thus, when isolating a single pair of positions, epistasis predictions are mainly informed by the stability of the individual mutations in another manifestation of learned assay-manifested global epistasis. A modified version of ESM-MSR (ESM-MSR masked marginal) that is trained to compute double mutant effects while concealing the mutated position identities shows the same type of pattern and reduced performance in direct epistasis ranking relative to ESM-MSR (Supplementary Table 2). A key difference between ESM-MSR and all other transfer learning methods is that the mutated sequences are encoded separately from the wild-type, requiring an additional full forward pass through the ESM3 backbone (with the MT LoRA adapter) per variant. This procedure allows the model to directly build representations based on the mutated sequence, while the other methods apply post-hoc mechanisms to insert the mutant-interaction information to representations derived from the wild-type sequence only. As evident from the good performance of the ESM-MSR masked marginal ablation, much of the variance in the data can be explained with identity-independent effects, and so the identity-dependent signal may be too small or too challenging to extract from the latent embeddings for most transfer learning approaches.

### Advantages and Limitations Based on Mutation Type

To gain insight into their practical utility, we probe the mutational preferences of the models and how different types of preferred mutations contribute to stabilization. Our retrospective testing strategy involves selecting 96 top-scoring single substitution mutants per domain from the Megascale test set and then assessing how those preferred mutations distribute into feature-based categories and contribute stability, visually described in Figure 3a. This experimental outcome of interest is the expected total stabilization achievable across all selected mutants during a medium-throughput screening experiment conducted in a 96-well plate. The results are shown in Figure 3b for classes based on physicochemical characteristics; the number of selected mutants in each class (lower subplot of b) inform a model’s category preference, while the rectified gain (upper subplots) inform the stabilization conferred by selected mutations from that category. We also introduce and explore the *σ* parameter, which is a scaling factor that controls the extent to which ESM-MSR’s predictions can deviate from the original predictions by scaling both LoRA alpha parameters relative to their original value (*α*_*WT*_ = 4*σ, α*_*MT*_ = 16*σ* ) .

**Figure 3.**
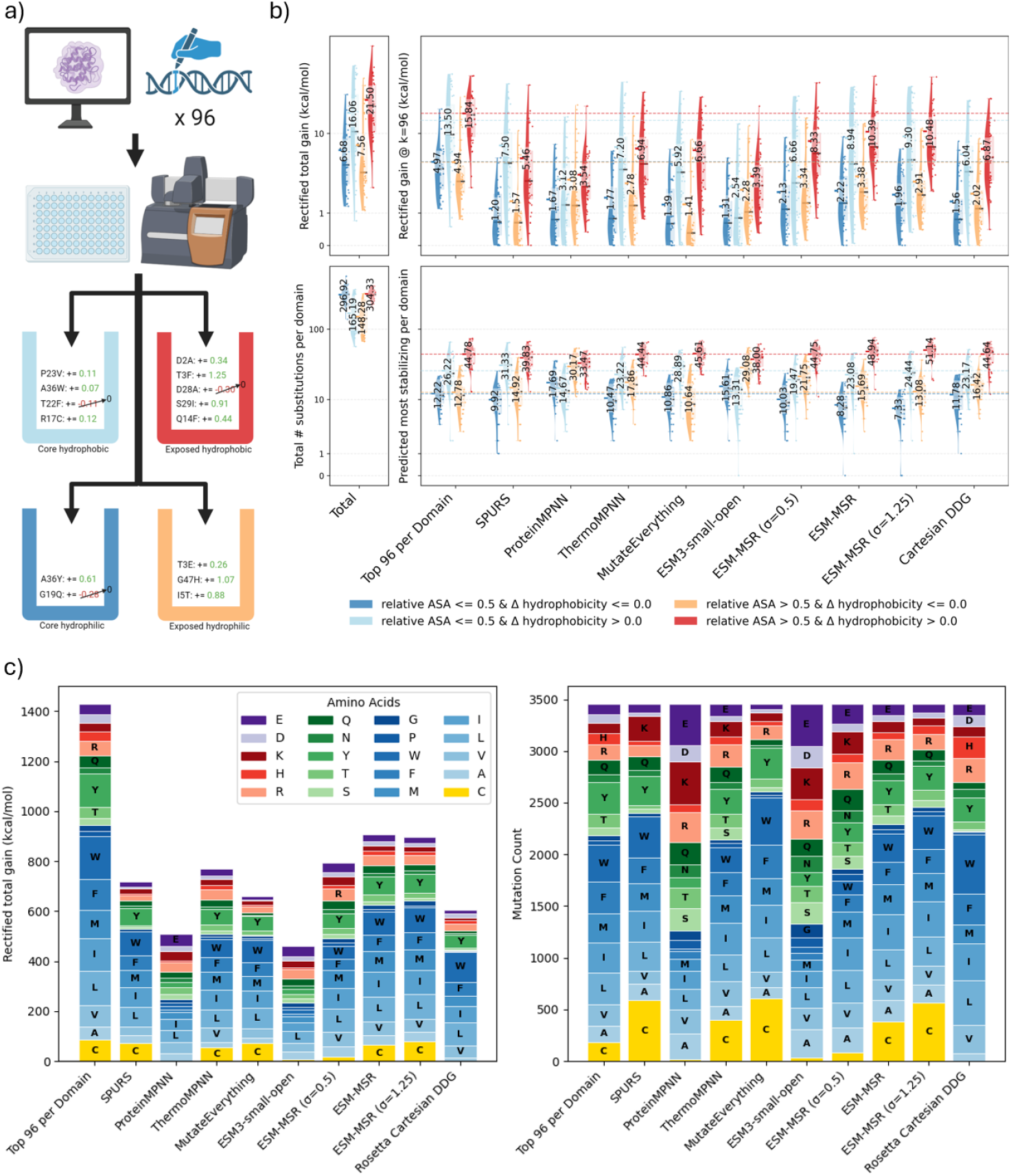
Retrospective Analysis of Top Scoring Substitutions from the Megascale Test Set. a): Schematic of the hypothetical experimental setup whose outcome is shown b). Each model selects its 96 top-scoring single substitution mutations for retrospective “experimental validation”. Positive stabilizing effects from mutations are added together, while destabilizing mutations are set to zero (but are still counted towards model preference). This process is repeated for each of the 36 test domains and averaged. b): Physicochemical Characteristics of Selected Mutants. Selected mutations are divided into one of four categories based on relative solvent accessible surface area of the mutated position in the wild-type structure (relative ASA, with 0.5 indicating that 50% of the wild-type side chain area is solvent-exposed) and change in Kyte-Doolittle hydrophobicity upon mutation (Δ hydrophobicity, negative values imply increased hydrophilicity). Each point represents a single test set domain; the upper subplot shows the combined stabilizing effect of all selected mutations in that category and the lower subplot indicates the number of selected mutants for that category. The points collectively form a distribution over all test domains, whose median is shown with a thick horizontal black dash and inner quartiles are indicated with a shaded box. Mean values are annotated with text and a colored dash. All values are displayed on a log y-axis to facilitate comparison. Leftmost panels indicate the full distribution of mutations, without selecting just the top 96. “Top 96 per Domain” is a positive control indicating the true experimentally most stabilizing substitutions per domain. Horizontal dashed lines colored according to categories represent the mean across domains for this control, to facilitate best-case comparison. c) Similar to b), except that all selected mutants are pooled together to assess the total gain conferred by specific mutant amino acid identities (left) and the mutation preferences (right).

In each of the four physicochemical property scaffolds, ESM-MSR outperforms all other methods in terms of achieving stabilizing effects in selected mutants, and cutting the LoRA strength in half (*σ* = 0.5) results in submaximal performance only relative to ThermoMPNN and SPURS in the buried hydrophobic category. Furthermore, except in the buried hydrophilic category, ESM-MSR obtains more than 65% of the average stabilization obtained from the experimental top 96 most stabilizing substitutions (referred to henceforth as the “best-case”). All supervised methods choose far more hydrophobicity-increasing, solvent-exposed mutations than any other category, while ProteinMPNN and ESM3-small-open choose relatively fewer and gain less than a quarter of the best-case stability. Hydrophobic surface mutations empirically offer the largest potential stabilization (Figure 3b, upper left panel), but these may reduce soluble expression when they cannot be buried or lead to difficult-to-predict structural rearrangements impacting function [14]. Instead, zero-shot models have a higher relative preference for hydrophilic mutations near the surface, matching previous results [37] and supporting the notion that models of natural sequence likelihood must compromise between stability and solubility to reflect natural fitness. Tuning the *σ* parameter from 0 (equivalent to ESM3-small-open) to 1.25 mainly results in an increasing preference for hydrophobic residues over hydrophilic ones, meaning the parameter provides some control over selection bias. Later, we show how this parameter can be used to optimize mutant abundance beyond even the level of ESM3.

Considering that they model natural sequence likelihood, it is interesting to compare the specific mutant residue identity preferences learned by supervised models relative to the baseline of zero-shot predictors. For instance, an unusual consequence of training on the Megascale dataset is a measurement bias that causes cysteine mutations to appear extra stabilizing. This bias results from surface-accessible cysteines forming intermolecular disulfide bonds, inhibiting proteolysis (the measured effect) without actually improving thermodynamic stability (the inferred property) [45]. Figure 3c shows that such mutations comprise less than 10% of the best-case selection pool in terms of both population (right panel) and stabilizing effect. All transfer learning methods over-select this category, especially SPURS and MutateEverything. This is an undesirable behavior that is not replicated in ProteinMPNN, ESM-small-open, or Rosetta, but reducing ESM-MSR’s *σ* to 0.5 almost completely eliminates the bias. Adding detail to the selection described in 3b, the best-case breakdown in c) shows that bulky hydrophobic amino acids, cysteine, and tyrosine are frequently among the most stabilizing possible mutations, and all transfer learning methods clearly reproduce this pattern. Indeed, scaling the *σ* parameter from 0 to 1 predictably shifts the distribution from the zero-shot ESM3 to a distribution that strongly resembles the best-case selection, but *σ* = 1.25 begins to distort the distribution. Zero-shot models are much more likely to select charged amino acids and all types of polar amino acids, explaining their relative hydrophilicity-increasing preference which better captures natural amino acid frequencies. Although partly trained to reproduce amino acid preferences, Rosetta shows the greatest preference for bulky hydrophobic residues among all methods, revealing a bias in the force-field parameters.

We extended our analysis to double mutants in Figure 4, where we chose to screen all possible residue pairs (over 16 million mutants) rather than attempt to draw conclusions from the 2-4 positions per protein experimentally assessed in the test set (10,732 mutants). A plot of epistasis versus minimum wild-type side-chain heavy atom distance (4a) shows that ESM-MSR is strongly biased to predict that any double mutant will be more stabilizing than the sum of individual effects, compared to the more symmetric ESM3. In either case, the vast majority of pairs (especially distant ones) are predicted to have epistasis very close to zero. ESM-MSR’s overall bias, which sometimes extends to predict positive epistasis for even distal mutants, is a manifestation of the global epistasis and survivorship bias explained previously. Higher-magnitude interactions are broadly distributed below a wild-type heavy atom distance of 10Å where larger mutant residues can form intermolecular interactions even if the wild-type residues are too distant. It should be noted that characterized double mutants in the Megascale dataset are heavily biased towards positions physically near enough to interact in the wild-type protein domain, exaggerating the importance of non-additive effects relative to unbiased positions, which tend to be non-interacting and near-additive [8]. Thus, it is encouraging that the magnitude of the ESM-MSR-predicted epistasis diminishes and approaches zero for increasingly distant mutant pairs, suggesting that ESM-MSR captures the expected relationship and is not undermined by this training data bias.

**Figure 4.**
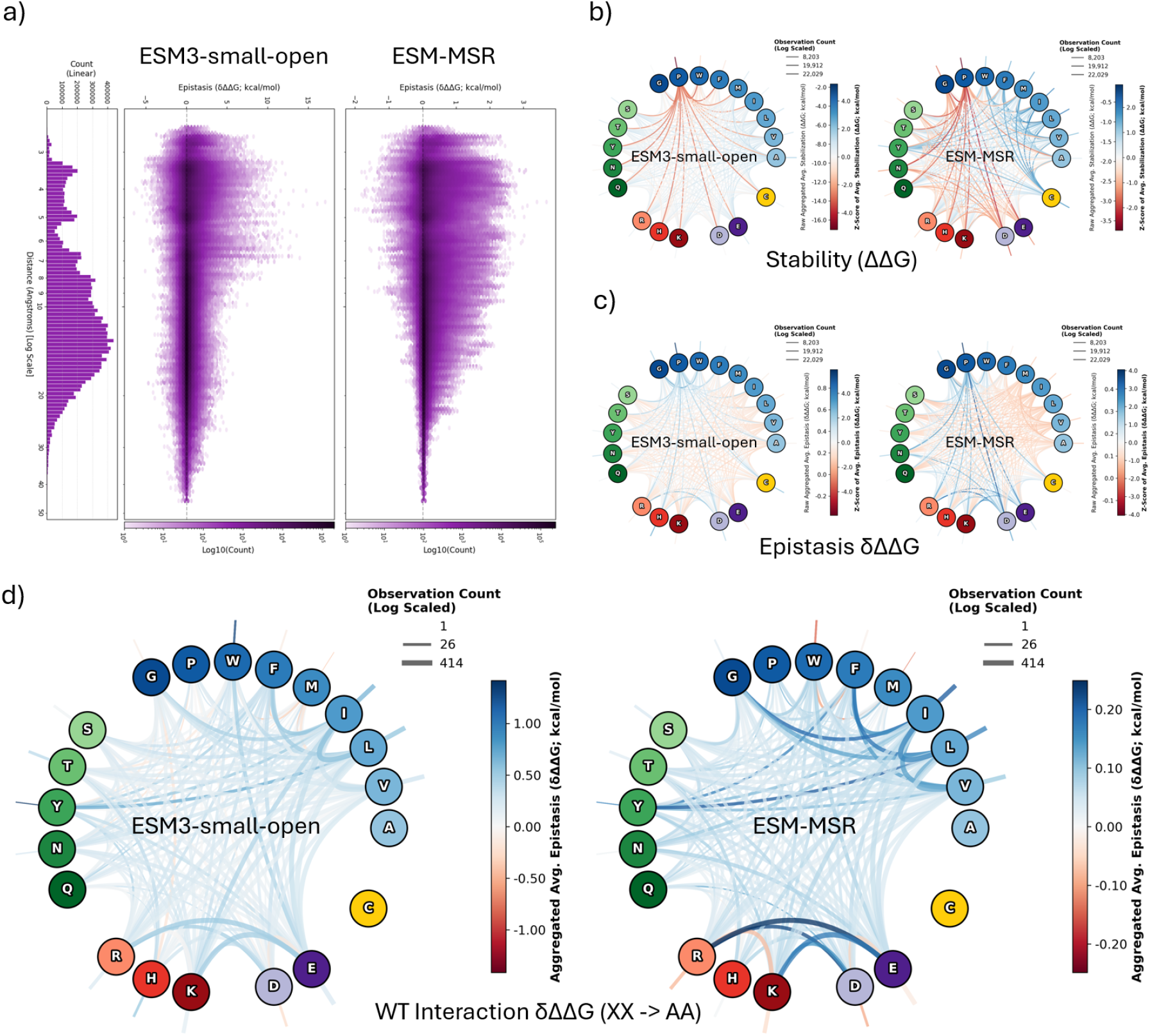
Double Mutant Screening of Test Set Domains. a): Summary of epistatic interactions when using ESM-MSR to screen all pairs of mutations, pooled from all 36 test set domains. Each hexagon represents a distance-epistasis bin, with darker purples indicating exponentially higher populations. b)-d) Pairs are filtered to those with a minimum side-chain heavy atom distance below 6 Angstroms. b): Average ΔΔ*G* predictions for each pair of residue identities (upper: wild-type identities; lower: mutant identities). Amino acids are represented as nodes, grouped and colored by their properties. The connections between nodes indicate the normalized, mean-aggregated stability property of mutants comprised of the connected node identities from the screen. Thicker and more darkly colored edges represent larger-magnitude z-scores, with blue indicating higher stability and red indicating lower stability. Exterior line segments represent double mutants to the same residue. c) Same as b), but now showing predicted epistasis. d) Same as b) and c), but filtered such that only double mutation to alanine is included to search for wild-type interactions. Normalization is removed to highlight emphasize raw epistasis scores, which also have very different magnitudes between the models.

The strongest predicted positive epistasis tends to occur via compensatory mechanisms (the range between D and E in Figure 1b): for instance, the top two points for both models involve mutation of a leucine-alanine pair in the core of 1UFM to glycine and either tyrosine or phenylalanine. Although the glycine mutation may cause a core-collapse on its own and the aromatic one would clash with the extant leucine, in combination, they roughly occupy the original volume. The introduction of opposite charges in hydrophobic environments also commonly leads to very positive epistasis and is explained by the electrostatic compensatory gain of a putative salt bridge. Meanwhile, the two most negative epistasis examples are seen in the triple helix *de novo* design HHH_rd1_0516, where a pair of proximal lysines are mutated into positively charged residues; each single mutant eliminates repulsion, but the combination restores it. Introduction of a same-charge pair into a clashing environment is a common motif among the most negative predicted epistasis examples, matching theoretical expectations.

To focus on direct interactions, we next filtered our screen to select pairs with a minimum wild-type side-chain heavy atom distance of 6 Angstroms or less. Comparing the left side panels of Figure 4b, ESM-MSR shows an increased overall preference for creating hydrophobic pairs versus ESM3, while possible salt-bridge and hydrogen bond-forming pairs are disfavored on average, likely due to their specificity requirement. Double mutations to proline are predicted to be most destabilizing, but all mutations involving proline tend to score lower than any others, regardless of which model is used. Indeed, proline-involving double mutants are so destabilizing that they appear to trigger the ESM-MSR’s learned assay floor, exhibiting high positive epistasis (Figure 4c). This unwanted behavior is less apparent in the zero-shot predictions from ESM3, but surprisingly is still present. Aside from this pattern, the epistasis predicted by ESM-MSR seems to reinforce (and recalibrate) the patterns seen in ESM3 with high positive epistasis consistently occurring between oppositely charged residues, but with newly emphasized epistasis between polar and charged residues.

Leveraging experimental biochemistry methodology, we filtered our data to exclusively alanine double mutants to isolate the wild-type contribution to epistasis and look for putative interacting residues in Figure 4d. In a simplified view, mutating either partner to alanine will likely break their interaction, while the second alanine mutation is unlikely to have a significant further effect. Thus, the stability epistasis approximates the interaction energy between the wild-type residues, *I*_*AB*_:

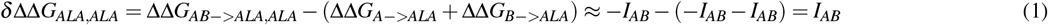

The two plots show that ESM-MSR is much more sensitive to identity-based interaction clues than ESM3, with high Z-scores for nearby oppositely charged residues and many hydrophobic pairs (including tyrosine), but low Z-scores for very bulky or same-charge pairs. Overall, while these observations demonstrate that ESM-MSR learns and prioritizes stability-relevant aspects of amino acid interactions, it also highlights the need for careful interpretation of the epistasis predictions of ESM-MSR in light of the assay limitations of its training data.

### Generalization to Full-Sized Proteins

Although ESM-MSR achieves superior ranking performance and theoretical low-throughput capitalization when testing on the held-out scaffold of the Tsuboyama Megascale dataset, this scaffold has key characteristics in common with the training data: both use protease susceptibility as a proxy measurement for stability and include only small protein domains (40–72 amino acids). To assess generalization toward practical applications, we first use thermodynamic measurements of full-size proteins, where previous methods have failed to generalize [5]. We note that the train/validation/test split used throughout this work is designed to minimize sequence identity between all training and testing data, including all external datasets described in this section, to minimize data leakage (see Supplementary Figure 1). Reliably benchmarking and comparing supervised models on external thermostability datasets is challenging, especially because data leakage control methodologies can vary widely between studies. In this study, we have controlled data leakage strictly while retaining as much training data as possible, and retrained as many methods as practical on these splits. For all methods except SPURS, we were able to implement the respective authors’ training workflows to produce models with comparable benchmark results to the original papers. However, we were unable to reproduce benchmark results when retraining SPURS, so instead we used their provided, pretrained model. We note that this model almost certainly leaks data, especially since the original authors did not hold out datasets such as K2369 in their study. Hence, SPURS results may be overly optimistic. After matching training splits, we tested the majority of recent high-performing stability models on a comprehensive set of thermostability benchmarks.

Figure 5a summarizes the performance of all tested models on heterogeneous datasets of experimental stability measurements of natural protein variants; ΔΔ*G*, the free energy change upon unfolding is used except for S571 which uses Δ*T*_*m*_, the change in melting temperature. ESM-MSR demonstrates the highest average performance, to our knowledge achieving state-of-the-art performance on four of the ten benchmarks: S461, S783, K2369, and PTMUL-D. The second most robust model is MutateEverything trained *without* double mutations, since we found that training on these mutants adversely affects performance on the single-mutant test sets. ‘WT’ approximations generated by ESM-MSR (white triangles) provide excellent performance on single mutants by running a single wild-type sequence forward pass through only the WT path of the model per protein (see Figure 1a). Unexpectedly, averaging-in the MT LoRA path (the ‘mutant’ perspective) slightly improves performance for singles, albeit at the high cost of switching from O(1) efficiency to O(19*L + 1) (1 pass per mutant plus the wild-type sequence), perhaps by more strongly encoding the possible interactions of the mutant residue. The addition of the MT path is much more impactful for multi-mutants, consistent with the theory that it helps the model to encode epistasis through directly encoding mutant identities in the input. However, it is also expensive for comprehensive screens (see Supplementary Table 4&5 for complexity and inference cost analysis). Also surprising is the continued improvement in ranking scores that is often seen when increasing *σ* to 1.25. As shown in the previous section, the higher LoRA alpha values may allow ESM-MSR to further shift its mutational preferences to better exploit the biases of thermostability benchmarks, especially toward surface hydrophobic substitutions. In terms of destabilization bias, Supplementary Figure 10 shows that ESM-MSR and SPURS achieve the best direct-inverse antisymmetry on Ssym (-0.87, ideally -1) and ESM-MSR is the least biased (-0.160, ideally 0) among tested models despite a lack of auxiliary losses or data augmentations enforcing these behaviors during training.

**Figure 5.**
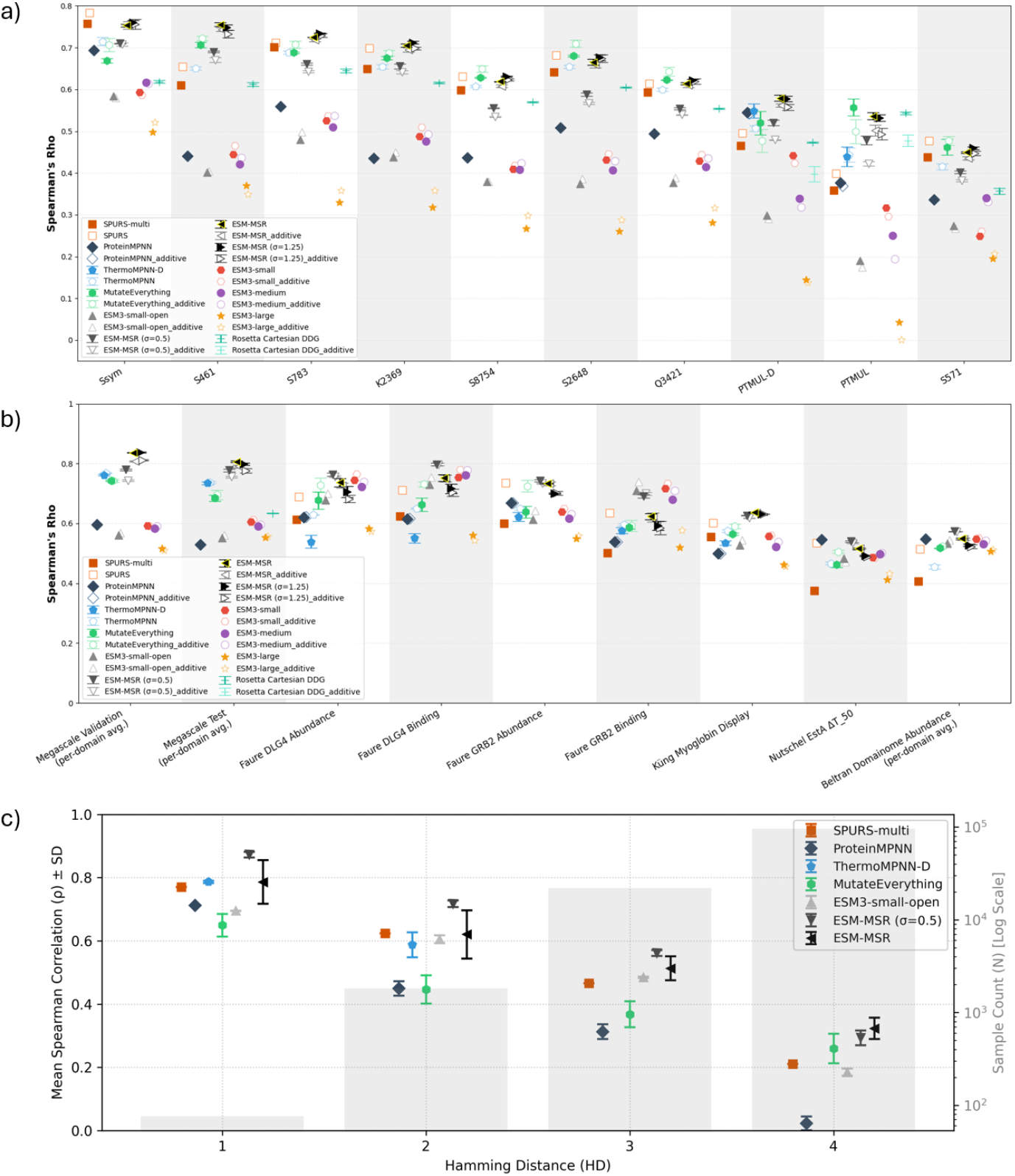
Generalization of Stability Predictors to External Datasets. a): Average ranking performance on benchmark thermostability (ΔΔ*G*) datasets. Error bars represent one standard deviation around the mean computed from 3 replicate models trained on the same subset of the Megascale dataset with different seeds (except for SPURS, which was originally trained on a different subset). For Rosetta, the replicates are from 3 pairs of wild-type and mutant relaxed structure predictions. Markers for each model represent alternative inference strategies. For ThermoMPNN, filled markers use a combination of ThermoMPNN-D for double mutants and ThermoMPNN predictions single- and (additive) multi-mutants, since ThermoMPNN-D can only predict double mutants. White markers indicate that only additive predictions from ThermoMPNN are used. For SPURS and MutateEverything, we observed that models trained only on single mutants performed best prior to additional training on double mutants, so white markers indicate singles-only training and additive multi-mutant prediction while filled markers represent the fully trained models (SPURS-multi and MutateEverything) used natively. For all ESM3-based models (including ESM-MSR, which is always trained with single and double mutants) white markers represent only performing a WT LoRA pass using one sequence per protein, scoring additively for multi-mutants, with the full two pass “dual-view” combined average for filled markers (see Methods). b): Average ranking performance on other properties, each explained in the text and Methods. The Domainome abundance dataset points represent the mean Spearman’s *ρ* for ranking the relative abundance of mutants across 522 human protein domains, while the Megascale benchmarks are identical to those from previous sections with SPURS removed due to train-test overlap. Error bars and outline-only markers have the same meaning as in a). c) Performance by mutation count on the GB1 epistatic fitness landscape. Spearman *ρ* correlations are for mutants grouped by Hamming distance to the wild-type sequence. Shaded bars indicate the number of mutants on a log scale. Since the focus is epistasis prediction, additive ThermoMPNN predictions are not shown for higher order mutants.

ProTherm-Multiple (PTMUL) and its double mutant subset (PTMUL-D) are of particular interest for assessing generalization, representing the most comprehensive datasets of multi-(double-) mutant thermostability effects, containing 912 (536) unique mutants with up to 10 (2) mutations [27]. ESM-MSR exceeds the ranking performance of all methods tested in PTMUL-D. All methods except SPURS improve on their respective additive baseline, contradicting Figure 1d, in which only ESM-MSR improved on the Megascale test set. Meanwhile, both methods incorporating ProteinMPNN representations (SPURS and ThermoMPNN-D) fail to significantly improve over the zero-shot approach. MutateEverything and Rosetta slightly outperform ESM-MSR on the full PTMUL set including multi-mutants, but not on the double mutant subset. With each additional mutation added to a protein, the probability of structural arrangement increases significantly. Through its geometric attention and structure-token modalities, ESM-MSR is heavily reliant on the structural input, but for multi-mutants, this structure is likely inaccurate. Hence, we attribute MutateEverything’s better performance to its more appropriate inductive biases, obtained by conditioning on the multiple sequence alignment and only implicitly inferring structural elements from evolutionary context. Meanwhile, Rosetta is the only method studied that attempts to predict the mutated structure, a critical addition for multi-mutant modeling [33]. However, Rosetta’s strong performance is confounded by the use of ProTherm mutants for validation while developing Cartesian DDG [15].

### Generalization and Trade-Offs with Other Properties

Self-supervised protein models such as ESM3 have repeatedly shown excellent zero-shot predictive performance for mutant effects on properties other than stability [29]. This ability highlights the connection between the likelihood of observing a mutation in nature (the modeling objective) and protein fitness, with the combined effects of many properties (e.g. stability, expression level, and biological activity) contributing. ESM-MSR provides an opportunity to directly explore the trade-off between zero-shot predictions, correlated with multiple aspects of fitness, and stability-specific predictions, which may compromise soluble expression (cellular abundance) or function (e.g. binding to a functional partner). Specifically, we can scale ESM-MSR’s *σ* from 0 (equivalent to ESM3-small-open) to 1 (ESM-MSR) and beyond, where an intermediate value of 0.5 represents a compromise between stability and general fitness. Such tuning is not easily achieved with other approaches like Mutate Everything and ThermoMPNN, which operate on latent embeddings. While setting *σ* to 0.5 consistently reduces the ability of ESM-MSR to predict the effects of mutations on the unfolding free energy change (Figure 5a), the deep mutational scanning (DMS) assays in Figure 5b show that generalization to other properties is usually improved. Each comprises thousands of experimental measurements of mutant effects on properties other than thermodynamic stability, and *σ* = 0.5 tends to perform best.

First, a study from Faure and colleagues [12] considered binding and cellular abundance of single- and double-mutants of two common human interaction domains: GRB2-SH3 (binding to GAB2) and PSD95-PDZ (DLG4, binding to CRIPT). The cellular abundance of protein variants correlates with their thermodynamic folding stability because destabilized mutants populate unfolded or misfolded states more frequently, leading to increased engagement with quality-control pathways and proteolytic degradation. Among the four assays, we see large improvements through fine-tuning, with ESM-MSR (*σ* = 0.5) significantly outperforming ESM3 except on GRB2-SH3 binding, where it performs similarly. Binding generally involves a biophysical mechanism in which free energy is reduced through interaction with a binding partner, often favoring conformational flexibility or marginal stability. ESM-MSR has necessarily traded-off its understanding of some aspects of fitness to prioritize stability prediction, although lowering the LoRA’s influence compensates or even reverses this behavior. This result contrasts the behavior for predictions corresponding to a study from Küng et al. [21], who measured yeast cell surface display for single- and double-mutants of human myoglobin, expected to correlate strongly with mutant stability and expression. Here, the default ESM-MSR (*σ* = 1) outperformed all other methods, including the base model. In the EstA DMS experiment from Nutschel et al., change in *T*_50_ (the temperature at which an enzyme variant has lost 50% of its original activity, relative to the wild-type) was assessed for single mutants of Lipase A from *Bacillus subtilis* [30]. ESM-MSR improves over the zero-shot baseline (ESM-small-open), but *σ* = 0.5 further enhances performance, matching the zero-shot performance from ProteinMPNN. Notably, ESM-MSR appears to retain or very slightly increase in performance when considering epistasis in the above datasets, while all other transfer learning methods tend to perform much worse when attempting to infer epistasis. In general, additive ThermoMPNN has much better performance than ThermoMPNN-D and inconsistent improvement over its ProteinMPNN base model, unexpectedly gaining performance on binding benchmarks while ranking score decreases on Δ*T*_50_ and most abundance assays. ThermoMPNN uses a more data-dependent form of transfer learning than the low-rank adaptation used by ESM-MSR, which may lead to greater variance on test sets.

As a rigorous test of generalization, we examined a comprehensive study from Beltran and colleagues, who measured single-mutant abundance data for 522 human protein domains [2]. Across 536,145 mutations in this massive dataset, the highest-performing approach is ESM-MSR (*σ* = 0.5), achieving an average Spearman’s *ρ* of 0.573 and slightly outperforming both ESM3 and ProteinMPNN. ThermoMPNN, which was the highest-performing predictor assessed in the original study by Beltran et al., performed significantly worse than all others, including its own zero-shot baseline, when trained on our Megascale training subset. This benchmark provides conclusive evidence that ESM-MSR effectively models stability without sacrificing abundance, as the full *σ* = 1 model performs slightly better than the zero-shot ESM3 base model. Notably, the computational cost of 536,145 predictions is considerable for our dual-view approach, requiring approximately an hour to complete on an RTX 5090 GPU. However, since the benchmark is comprised of only single mutants and the performance using the WT approximation is virtually identical, this fast heuristic is recommended for high-throughput single mutant screens. Evaluating the full Domainome requires just 522 forward passes with this strategy, effectively a 1000x speed-up over evaluating each mutation.

Finally, we explored a formidable challenge: predicting the near-exhaustive quadruple mutant fitness landscape at four interacting positions of GB1 [46], shown in Figure 5c. Naturally, all methods see substantial, near linear degradation in ranking performance with mutation count, with ESM-MSR (*σ* = 0.5) clearly leading except for quadruple mutants. Interestingly, Mutate Everything’s degradation appears more gradual, perhaps because it does not rely on increasingly inaccurate structural inputs. This behavior, also seen in PTMUL, reinforces the idea that structure-dependent models are limited by their inputs, possibly suggesting a gap for accurate mutant structure predictors to fill. By contrast, ProteinMPNN relies almost exclusively on structural clues (backbone coordinates) and shows the greatest performance degradation for multi-mutants, while ESM-MSR, ESM3 and SPURS rely both on structure and sequence. Overall, ESM-MSR (*σ* = 0.5) exhibits unexpected high relative performance even for multi-mutants on non-stability properties, but improvement over the zero-shot approach is inconsistent in comparison to abundance.

## Discussion

Although they are often strong zero-shot predictors of the effect of mutations on fitness-associated protein properties, self-supervised protein models face limitations in stability prediction arising from their original training data and objectives. Transfer learning can specialize these pre-trained models, overcoming these limitations by providing a strong and direct stability signal and examples of diverse stabilizing mutations absent during pre-training. In this work, we demonstrated a new transfer learning approach, ESM-MSR, with capabilities that exceed contemporary stability predictors in two key areas: selecting stabilizing mutations and scoring epistatic double-mutants. These advances were likely enabled by a key difference between our approach and previous methods: we fine-tune the base model end-to-end in a near-native inference mode, applying LoRA to ESM3 in order to adapt differences between mutant and wild-type residue logits into thermostability predictions, aided by a basic linear calibration head. Our approach contrasts SPURS, ThermoMPNN and Mutate Everything, which must learn data-intensive transformations from latent embeddings to physical stability quantities. In other words, ESM-MSR has a “head start” for data-efficient training, exploiting the connection between stability and residue likelihood as an inductive bias.

An ongoing challenge in protein engineering is to accurately identify stabilizing mutations without adversely affecting (or ideally, while improving) other properties, especially solubility. Researchers have repeatedly demonstrated that learned biases limit the practical applicability of stability models, especially since they predict hydrophobic surface mutations as more stabilizing and most other types as more destabilizing than the true distribution [10, 4, 34, 40]. We established the ability of ESM-MSR to select the most stabilizing mutants across physicochemical property scaffolds and to rank and retrieve stabilizing mutations in our Megascale test set. Additionally, we established state-of-the-art ranking performance on multiple thermostability benchmarks without reducing ranking performance on abundance assays relative to the zero-shot prediction. Although we note that ESM-MSR still inherits biases that could result in high scores for cysteine mutants and mutations that can reduce solubility or harm natural function, we demonstrated that the *σ* parameter can function to mitigate these biases. More generally, it can act as a tool for exploring the stability-function trade-off: we found that *σ* = 0.5 surprisingly improved abundance-ranking performance over both the zero-shot and tuned models among 522 assays in the Human Domainome. Based on the amino acid preferences shown in Figure 3b, it seems most plausible that this improvement is achieved by contextually shifting likelihood distributions based on stability without fully deviating from the natural residue distribution, which contains more polar and charged residues, preserving both the solubility and stability required for abundance. Although ESM-MSR (*σ* = 0.5) sacrifices some stability performance to achieve this, it still far exceeds ESM3 and competes with weaker stability predictors including Rosetta, making it an interesting model for protein engineering applications. In particular, the low *σ* setting will likely be useful for rescuing heterologous expression by improving both folding stability and solubility. In future work, it will be important to study the trade-off in enzyme activity resulting from increasing *σ*, especially since enzymes tend to rely on low-stability active sites to promote substrate binding and chemical reactions. In the absence of such understanding, the popular PROSS tool has seen practical success simply by restricting mutagenesis to non-conserved sites before applying Rosetta single-mutant prediction [16]. Labs with limited wet-lab resources could apply the same paradigm with ESM-MSR as the predictor and extend screening to double mutants to achieve low-N experimental stabilization of natural enzymes.

Unreliable prediction of non-additive multi-mutant effects is a significant barrier in protein science that restricts the mutational space that can be effectively screened *in silico*, necessitating expensive *in vitro* approaches to obtain stable multi-mutants. In this work, we demonstrated that ESM-MSR has explicit understanding of epistasis in addition to superior performance on double and multi-mutants from held-out test sets. However, we also showed that all methods trained on the Megascale dataset are essentially incapable of ranking negative epistatic effects. This failure mode is symptomatic of a fundamental assay limitation: any sufficiently destabilized mutant cannot be adequately characterized, and examples of negative epistasis are extremely rare since they exaggerate destabilizing effects. In order to provide this missing slice of training data, future assays could explore double mutagenesis of extremely stable *de novo* scaffolds. We also found that training ESM-MSR on only single mutants had no substantial impact on test set double mutant stability prediction, slightly degrading direct epistasis prediction and PTMUL-D ranking. Clearly, ESM-MSR can learn most aspects of epistasis without training on double mutants, which encode assay limitations that interfere with learning genuine assessment of biophysical interactions. Unlike Mutate Everything, ESM-MSR showed a degree of resistance to strong survivorship bias in the training set, but care must be taken when attempting to infer genuine, local, pair-identity-dependent interactions effects from all models trained on this dataset. We expect that researchers will find ESM-MSR more effective for screening applications, such as selecting initial stabilized mutants for subsequent activity engineering, than for specific mutant analysis, such as deciding which single mutants identified in directed evolution can be combined without clashing.

Numerous methodological limitations still restrict accurate multi-mutant stability prediction independently from the training data. As mutations accumulate, the protein structure is expected to increasingly deviate from the wild-type structure that was used to make mutant-effect predictions, which increasingly necessitates mutant structure prediction during inference. Furthermore, without reasoning about explicit side chain atoms (which none of the studied models do), models may struggle to predict the fine-grained physical interactions that contribute to specific aspects of protein stability such as hydrogen bonding. Protein flexibility and dynamics pose an even more formidable barrier; they must be assessed to achieve a complete picture of stability, yet they are very difficult to model and currently expensive to predict. However, new high-throughput experimental datasets may enable models to begin to address this concern in the near future [13]. Rosetta addresses each of these aspects to some degree, yet its single-mutant performance is consistently lower than the machine learning models that ignore all of these aspects. Addressing these limitations in machine learning models, possibly by integrating explicit biophysics or auxiliary structure models, will lead to further progress in this field.

The success of our relatively simple implementation suggests areas for further exploration in the fine-tuning of protein language models for variant effect prediction. Epistasis in language models is a nascent field of study [28] and to our knowledge, we are the first to fully exploit the prior understanding of epistasis in language models through supervised fine-tuning in native logit space (see Methods and Figure 2a). Our two-forward-pass dual-view formulation improves over traditional masked marginal scoring when both are fine-tuned, verified through our “ESM-MSR (masked marginal)” ablation (Supplementary Tables 1-3), and mitigates the identity-independent epistasis prediction behavior seen in SPURS and ThermoMPNN-D. However, this strategy does not appear to improve zero shot multi-mutant prediction, likely because it deviates significantly from the mask language modeling pre-training objective. Instead, a hybrid between these approaches (“independent masking”; see Methods) does improve over masked marginal scoring, and may generally improve zero-shot inference for all protein language models, albeit at considerably higher computational expense. This strategy can also boost inference efficiency over default unmasked prediction during multi-mutant screens; see Supplementary Table 4. Meanwhile, the paradigm of fine-tuning protein models in their native end-to-end configuration remains underexplored, especially in terms of how to apply LoRA. Our ablation experiments showed that including the query, key and value projections of ESM3 in fine-tuning barely improved performance (over a feed-forward network up/down projection-only ablation, ESM-MSR*, only FFN up/down), suggesting that the attention patterns learned by the base model are either satisfactory for stability prediction or susceptible to overfit. Attention patterns in protein language models are established to correlate with residue covariance and physical interaction [36], providing a suitable strong prior for our task. Furthermore, we found that convergence typically requires fewer than 5 epochs, and performance across benchmarks quickly saturates when increasing the WT LoRA rank above 1, suggesting that shifting ESM3’s internal representations to be more stability-focused does not require complex reprogramming. However, a deep exploration of the various approaches for applying LoRA and parameter-efficient fine-tuning in general was outside the scope of this work. We conclude that ESM3 is highly amenable to fine-tuning for stability prediction in the logit space by construction, and the same is likely true for other properties and protein models, especially those integrating sequence and structure.

## Methods

### ChimeraX GUI Tool

To facilitate the broad use of our methods, we developed a graphical user interface (GUI) for ESM-MSR that can be easily installed and used in ChimeraX molecular visualization software [25]. After a brief setup, the tool is designed to orchestrate mutational screening for a protein of interest and then visualize the results intuitively. In terms of screening capabilities, the tool allows users to screen all residues of a loaded chain, a selected subset, or pairs of nearby residues within a specified distance (to reduce double mutant compute expense). Users can specify the LoRA model to use (and whether to use the fast WT LoRA approximation), the masking strategy, and the sigma value, and the GUI automatically handles non-canonical residues and most PDB indexing issues.

In terms of visualization, extensive options are shown in Figure 6 below. From single mutant screens, users will be able to quickly identify the predicted most-stabilizing and most-destabilizing mutations, mutable backbone positions, and possible interactions with nearby residues (6a). From double mutant screens, they can visualize top-scoring double mutants (explicitly viewing the individual choices that contribute to the aggregated data seen in Figure 4b), predicted epistatic interactions between mutations (4c) or predicted wild-type interactions (4d, shown in 6b). Users may find the exclusion flags useful, such as omitting cysteine mutations due to the known cysteine stabilization bias of the fine-tuning data. Furthermore, positive and negative threshold enable one to focus on the predictions of interest, filtering out small-magnitude or destabilizing mutant effects from the visualization. Nearby residues and atoms can be shown and separately styled to facilitate rational design and interpretation. Different visual elements can have custom styling to reduce clutter, improve contrast, and generate informative figures.

**Figure 6.**
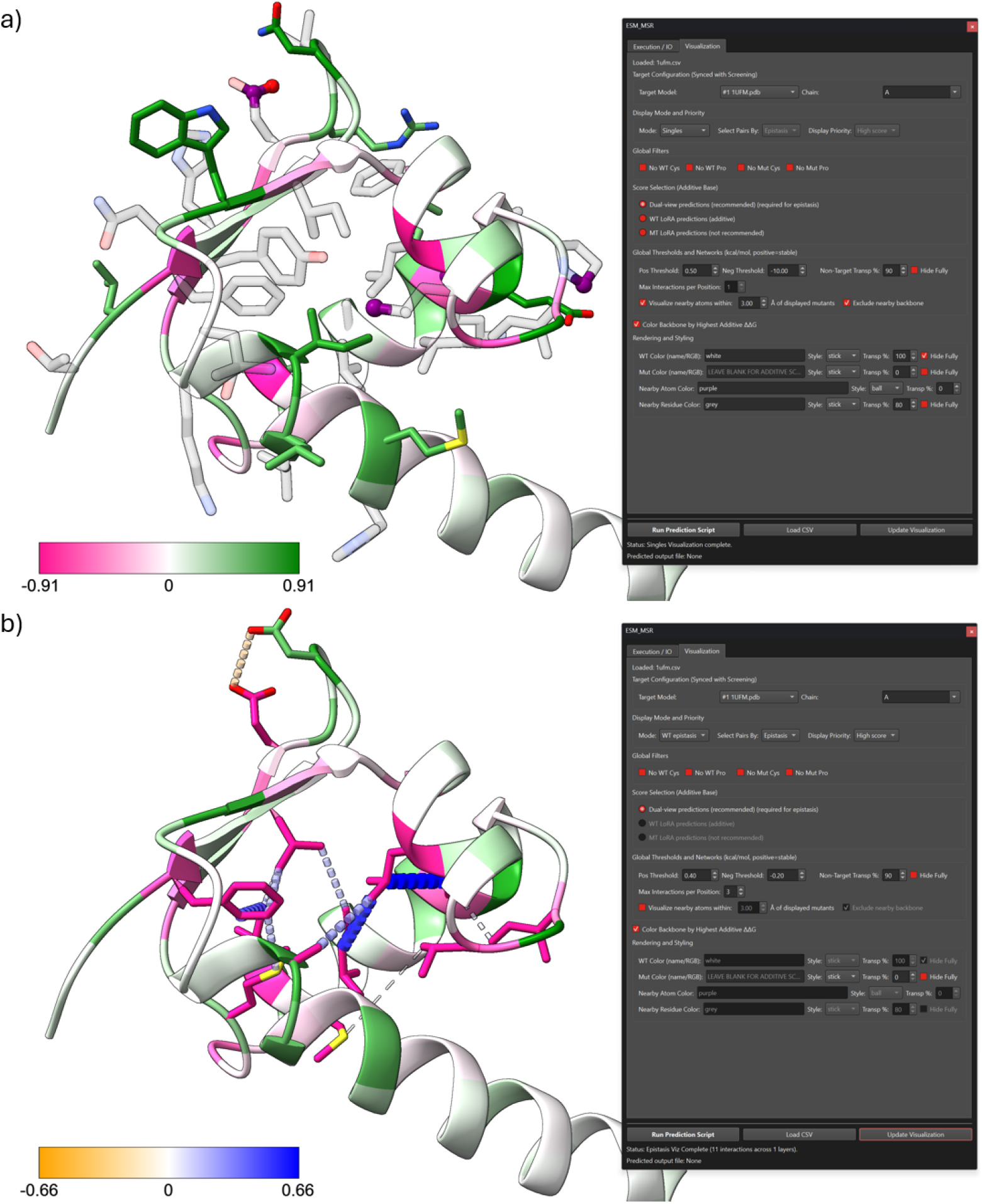
Screenshots of the ESM-MSR Tool in ChimeraX. a) Top single mutant screening candidates. With the displayed visualization configuration (right box), a full single-mutant screen is shown for the AlphaFold structure of 1UFM from the Megascale test set. The backbone is colored according to the most stabilizing residue at each position; pink positions are thus sensitive to any mutation while green positions may have opportunities for stabilization. The highest-scoring mutation at each position is visualized with green carbons if it exceeds the specified 0.5 kcal/mol threshold; destabilizing mutations would also be shown, but are filtered out by the low -10 kcal/mol destabilizing threshold. Nearby atoms to the displayed mutants (at a specified threshold of 3Å) are shown as purple balls, with the remainder of the residue shown in translucent grey. b) Wild-type epistasis network visualization. Wild-type residues are now displayed, with the coloring indicating the effect of mutation to alanine, shown as pink if these mutations are outside the range of the backbone coloring from a). Orange to blue pseudobonds indicate negative to positive epistasis and thickness indicates magnitude, but only effects outside the range [-0.2, 0.4] are shown, up to three per position. Epistasis is computed as in Eq.1, but with alanines additionally being mutated to glycine.

### ESM-MSR Implementation

#### Features

There are 7 possible inputs to ESM3 during inference, any of which can be disabled by masking. We input only the first three, which we believe to be the most salient for the prediction task: (a) sequence (amino acid tokens), (b) structure coordinates, and (c) structure tokens. The remaining input tracks ((d) 8-class secondary structure labels (SS8), (e) quantized solvent-accessible surface area (SASA) values, (f) function keyword tokens and (g) residue (InterPro) annotation binary features) are relevant to controlling generation, but are not expected to improve stability prediction and are omitted.

We refer the reader to the original ESM3 paper for a complete description of the features used by ESM3, and only summarize them here [17]. Regarding sequence (a), although ESM3’s vocabulary supports non-standard amino acids (B, U, Z, O), we canonicalize these amino acids when present during testing to match the restrictions of compared methods and because these non-canonical residues are absent from the fine-tuning data. The ESM3 with the WT LoRA adapter receives the tokenized wild-type sequence while the tokenized mutated sequence is sent to a copy of the model with the MT adapter. ESM3 only accesses the coordinates (b) of the backbone atoms: *C*_*α*_, *C*, and *N*. Consequently, the model cannot directly perceive side-chain atomic coordinates; it must infer side-chain packing and interactions from the amino acid identity provided in the sequence track. Unlike the discrete token inputs, structural coordinates are not embedded; instead, they are processed by a geometric attention layer integrated only into the first block of the transformer. Coordinates are padded on both sides with infinity, with no masking. Like the sequence tokens, structure tokens (c) are passed to the model flanked by beginning of sequence (BOS) and end of sequence (EOS) tokens without any masking. The structure tokens summarize local geometric context in a discrete format. These tokens are derived from the backbone coordinates using a VQ-VAE encoder that compresses the 3D information of the local neighborhood around each residue into a discrete code, providing a rich, learned representation of local shape without explicitly encoding side-chain coordinates.

#### Stability Predictions

We combine the wild-type marginal and mutant marginal formulations originally described by [24]. We calculate scores for each context as the sum of differences in raw logits assigned to the mutant versus the wild-type residue at each mutated position, separately conditioned on the wild-type and mutant sequences (Eq. 2 & 3). These scores are then calibrated and averaged between the two perspectives (Eq. 4). For the fast WT-only approximation (Eq. 5), we just skip the MT path and averaging, and the WT path is identical for every mutation of a given protein, so the logits only need to be computed once. We did not convert to log-likelihoods using a softmax over canonical identities since the result can be shown to be mathematically equivalent.

We define the stability prediction score Δ*LL* as the difference in logits assigned by the model to the mutant versus the wild-type residue at position *i*:

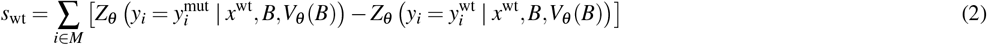

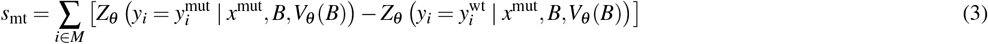

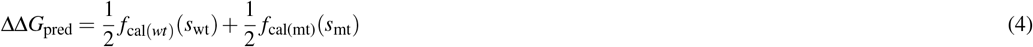

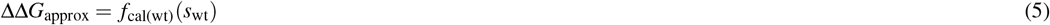

Where:

- *M* is the set of mutated positions.
- *x*^wt/mut^ is the full, unmasked wild-type / mutated tokenized sequence.
- 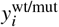 indicates the wild-type / mutant identities at position *i*.
- *Z*_*θ*_ is the model’s raw output logit for a specific residue identity at position *i*.
- *B* is the set of provided wild-type backbone atom coordinates.
- *V*_*θ*_ is the VQ-VAE that outputs contextual structure tokens.
- *f*_cal(wt/mt)_ is the learned linear calibration function.

The unmasked dual-view inference scheme has a considerable computational cost for protein-wide screening; although single mutants can be approximately scored in a single forward pass (ΔΔ*G*_*approx*_), using the MT path adds an additional L * 19 forward passes, where L is the protein sequence length. We used this latter dual-view score for single mutations (1 WT pass total and 1 MT pass per substitution) and compared this to the multi-mutant score when computing epistasis. Screening all possible double mutations requires 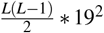 forward passes, which requires on the order of ten million forward passes, costing many GPU hours even with batching. This cost can be reduced 9.5 fold by iteratively masking each single position *i* in the sequence and running one WT forward pass (2) and one MT forward pass (3) in the masked context; “Independent Masking”, see Supplementary Table 4 for more details. Further, epistasis can be assumed to primarily occur between spatially proximal residues. Distal pairs of mutations can be computed rapidly using the WT approximation, and rigorous computation can be restricted to pairs within a minimum heavy atom distance, leading to roughly an additional order of magnitude compute savings. These efficiency approaches are optionally applied in our publicly available pipeline.

#### Low Rank Adaptation

We used the parameter-efficient fine-tuning Python library (peft) to apply LoRA to all transformer attention and fully connected layers in the full ESM3-small-open model, excluding the structure encoder. We used a rank-stabilized LoRA (RSLoRA) configuration with no bias, targeting only query, key and value projection “layernorm_qkv.1” layers (including the geometric attention in the first transformer block), and feed-forward network “ffn.1” and “ffn.3” up- and down-projection layers. Unless otherwise stated, we used a rank of 2 and an alpha of 4 for the WT LoRA (adding 2,073,728 parameters) and rank 16, alpha 16 for the MT LoRA (16,589,824 additional parameters). Combined, this represents less than 1.3% of the original 1.4 billion frozen parameters. We perform dropout for both LoRAs at probability of 0.1.

#### Training

We train the LoRA and calibration head parameters, with all ESM3 parameters frozen, using batches of mutants from one domain per batch. Each batch of mutants (*n* = 256) can contain any combination of single substitutions and double-mutant-derived mutated-context substitutions, and batches are constructed randomly, without repetition. Each batch of 256 mutants from a single domain is split into 16 minibatches of 16 mutants, and each minibatch is passed through the model to rank its mutants from most to least stable according to the equations above.

Computation is now split into two distinct graphs: the wild-type sequence, coordinates, and structure tokens are passed to ESM3 with the WT LoRA, while the mutant sequence and *wild-type* coordinates and structure tokens are passed to ESM3 with the MUT LoRA. For each “path” (WT and MUT), we compute the ListMLE loss of the uncalibrated logit scores *S* (explained below) for each minibatch to determine the gradient of the LoRA parameters, which are updated after each minibatch according to the AdamW optimizer. The label for the WT path is ΔΔ*G*_*additive*_, equal to ΔΔ*G* for single mutants or ΔΔ*G*_*A*_ + ΔΔ*G*_*B*_ for double mutants. The MT path applies teacher forcing, adding the reverse-calibrated ΔΔ*G*_*additive*_ label to *S*_*mut*_. The purpose of using uncalibrated predictions for the ListMLE loss is to stabilize the ranking loss by decoupling the prediction scale, and so reverse-calibration is necessary to rescale the additive label to match the logit space. If, for a given double mutation, both constituent mutation labels are not present, the predicted ΔΔ*G*_*additive*_ logit score is used instead. We also compute the gradient for the linear calibration head parameters based on the mean squared error loss of the calibrated output (ΔΔ*G*_*pred*_) to the label, which has a lower weight but also backpropagates into the LoRAs. This process is then repeated for a batch from the next domain, and mutations distribute into batches differently every epoch. We perform validation early stopping and evaluate the earliest model that achieved an average Spearman’s *ρ* score within 0.01 of the maximum.

#### Density Capping

To mitigate overfitting by exploiting the strong negative correlation between epistasis (derived *δ*ΔΔ*G*) and additive stability change (ΔΔ*G*_*additive*_), double mutations are binned into a 15×15 grid corresponding to these axes. Items from any 2D bin exceeding the 75th percentile among bin populations are randomly truncated down to that cap in each epoch, effectively flattening the typically sharp ridge observed on the best fit line between these properties. Double mutants without a derivable *δ*ΔΔ*G* label (because one or both single mutant measurements is missing) are capped to 20% of the population. Subsampling is repeated each epoch, allowing originally omitted samples to appear in later epochs.

#### Learning-to-Rank

We implemented the ListMLE loss function [47] based on the Plackett-Luce model. Unlike traditional regression objectives, which minimize the error between predicted scores and continuous experimental measurements, ListMLE maximizes the likelihood of the correct relative ordering of variants. This formulation (Eq. (6)) is theoretically effective for stability engineering as it prioritizes the correct ranking of the most stable variants (top-ranked items) over the precise ordering of unstable ones. The ground-truth permutation *π* is derived by sorting the experimental protease susceptibility measurements within each batch.

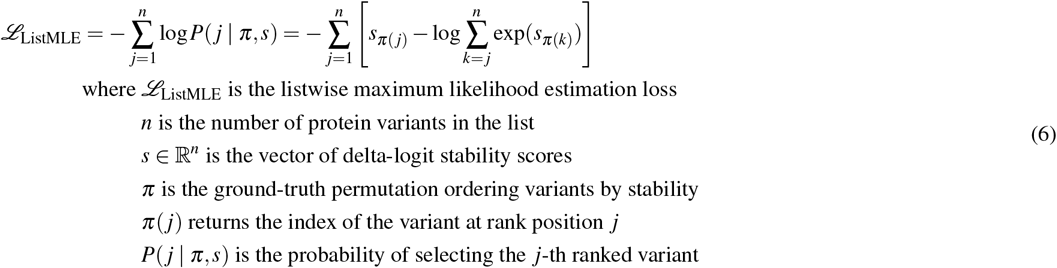

The ListMLE objective models the ranking process as sequential selection without replacement. For a batch of *n* = 16 variants, the probability of the true top-ranked variant (*π*(1)) is computed as a softmax over all n scores. This variant is then effectively removed from the pool, and the probability of the second-ranked variant (*π*(2)) is computed as a softmax over the remaining *n* −1 scores. This process repeats until the list is exhausted. By summing the negative log-likelihoods of these stepwise probabilities, the model is penalized heavily for misranking items at the top of the list. We compute this loss exactly over the full minibatch, rather than approximating it via subsampling as done in Zhou et al [50].

### Data

#### Preprocessing

Starting with the Megascale dataset table Tsuboyama2023_Dataset2_Dataset3_20230416.csv from Tsuboyama et al., we remove wild-type entries, insertions and deletions. For the same mutation represented by more than one sequence (e.g. when extra SAGG repeats appear before the target sequence), we take the average of the experimental measurements. We also remove “fake” mutations in which one or more of the constituent “mutations” listed is to the wild-type, and “improper” mutations where the same position is listed more than once with different mutations specified, which mainly occurs on destabilized backbones. The final dataset contains 521,154 mutations. Note that the “ΔΔ*G*” values from this dataset are estimated from raw read counts of cDNA-labeled domains determined by resilience to degradation by two orthogonal proteases. The final values are inferred based on an assumed two-state folding model [45].

Unlike the original implementations of ThermoMPNN and Mutate Everything, we train and test all models on mutants from structures with destabilized backbones. The original authors of the Megascale dataset applied destabilizing substitutions to highly stable domains to bring stabilizing mutant effects into the dynamic range of the protease susceptibility assay, since the assay depends on detectable changes in equilibrium folded fraction. For instance, the difference between a stable domain (population 99.9% folded) and a further stabilizing mutant (99.95% folded) was hard to reliably quantify. We found that training with these destabilized domains slightly improved performance for all models. We use the wild-type structure coordinates for these cases (rather than modeling the mutation), but edit the sequence to reflect the mutation. Since Rosetta requires all atomic coordinates (including side chains) and is the only tested approach capable of intrinsic mutant structure prediction, we reused the Cartesian DDG workflow to generate an initial structure including the destabilizing backbone mutation to score additional mutations using Rosetta. Hence, Rosetta may obtain a very small advantage over other methods for this class of mutations.

For Tsuboyama et al.’s Megascale dataset, we use the provided AlphaFold2 structures. In all cases, this includes only the mutated chain of interest (monomeric structures). The same is true for Human Domainome 1 structures, except that we obtained precomputed structures made available by the authors of SPURS [22]. No further preprocessing was done on the structures in either case. For deep mutational scanning datasets, including “Kung Myoglobin Display”, “Faure (GRB2 / DLG4) (abundance / binding)”, and Nutschel EstA dT_50, we generated AlphaFold 3 structures. Additionally details about these datasets is given in subsequent sections. For the GB1 epistasis datatset from Wu et al., we used the PDB structure 1PGA. Computing Rosetta Cartesian DDG predictions for the Domainome and DMS datasets was expected to take on the order of months, so these were not performed.

The following applies to external thermostability datasets: Q3421, K2369, S461, Ssym, S783, S2648, S8754, S571 and PTMUL(-D). In our preprocessing approach, we first download and preprocess the structures provided as PDB codes in the associated datasets. For NMR ensembles, we use the first model. We remove terminal capping groups, waters and heteroatoms before canonicalizing any nonstandard residues using PDBFixer’s replaceNonstandardResidues function. We also fill in any missing residues or atoms using Modeller (version 10.4). We note a small effect on performance on all methods caused by filling in missing residues, which may result in minor discrepancies with other reported results, but is probably leads to a more accurate structure. For S571, PTMUL and S8754, these sometimes include multiple measurements for the same mutations, for instance under different pH conditions. When this occurs, we simply average all measurements corresponding to the same mutant. This process reduces S571 to 551 unique mutants, PTMUL from 914 to 912, and S8754 to 6411. Note that S571 comprises Δ*T*_*m*_ measurements, while the remaining datasets are exclusively ΔΔ*G* measurements.

#### Train-Validate-Test Split

To create the main split used throughout the text, our primary concern was to mitigate data leakage into our external test sets. Hence, we filter at a maximum domain identity of 35% to any test protein in any test set; we refer to these as “low-test-identity domains”. We did not attempt to eliminate overlap with the 522 Domainome domains but found that only two domains had overlap with the training or validation sets.

First, all candidate protein sequences including external test sequences were clustered using MMseqs2. Any cluster containing one or more sequences from the external test set or exclusionary targets was strictly quarantined and removed from the candidate pool. The remaining “safe” clusters were randomly shuffled using a fixed seed to ensure deterministic reproducibility. To prevent intra-cluster leakage, clusters were assigned in their entirety to the validation set until a threshold of at least 30 proteins domains met, followed by population of the training set. These assigned sets retained all homologous members of the cluster. Finally, a rigorous post-assignment sweep was conducted utilizing O(N^2^) local pairwise alignments (Biopython, BLOSUM62) across partitions. Any sequence within the training or validation sets sharing sequence identity above a strict threshold of 35% with any testing protein was explicitly dropped. Crucially, we divide the number of alignment matches by the length of the small protein domain, effectively what fraction of that domain is contained in the full protein. While this filtration is very strict, it can still allow short motifs to leak. Validation sequences overlapping with the training set were also discarded, thereby guaranteeing rigorous sequence divergence across all evaluation boundaries.

In the main text, all methods are compared on an identical test set of 43,700 (32,968 single- and 10,732 double-) mutants from 36 small protein domains with low sequence identity to any training domain from the Tsuboyama Megascale dataset. This test set includes 2,302 mutations from domains where a single substitution has been made to destabilize the backbone to shift mutation effects within the detection range of the cDNA proteolysis assay. It also includes 25,593 mutations from *de novo* designed backbones. The latter subset may slightly advantage ESM-MSR over methods that depend on evolutionary information, especially MutateEverything. More detail is given in the Supplementary Information, Figure 9.

### Training and Testing Other Models

For Mutate Everything, we follow the original authors’ procedure as closely as possible using the code provided in their GitHub repository: https://github.com/jozhang97/Mutate Everything/. This involves generating multiple sequence alignments from the wild-type sequences using the ColabFold local workflow with default settings. For each of our dataset splits, we train for 20 epochs on single mutants only starting from the finetuning_ptm_2.pt checkpoint, and then 100 epochs with the full dataset, including single and double mutants. To the best of our knowledge, we exactly replicated of the original work, with the only deviation being the new training split(s) and our treatment of destabilized backbones, which involved correcting the input sequence. In this paper, we only consider the version of Mutate Everything built on the AlphaFold2 backbone, since it is shown to have superior performance compared to the ESM2-based version [31].

We train ThermoMPNN using the same AlphaFold structures used to train ESM-MSR. In this work, ThermoMPNN may refer to one or both of two distinct models used in a recent work [8]. Different models are used to predict single and double mutations, since each model can only predict one or the other. Double mutants use ThermoMPNN-D (a siamese neural network) and singles use the updated ThermoMPNN. Both are transfer learning models based on ProteinMPNN, and we use the implementations available from the GitHub repository: https://github.com/Kuhlman-Lab/ThermoMPNN-D. When we evaluate datasets containing double and single mutants, we combine predictions from both models. All higher-order mutant predictions are additive predictions from ThermoMPNN. PTMUL predictions use ThermoMPNN-D for double mutants and additive ThermoMPNN for the remainder. Again, we exactly replicated the original work except for the treatment of destabilized backbones, which were originally not considered or handled correctly.

We use the Rosetta Cartesian DDG procedure with the same settings used by Hoie et al. [19]. Our implementation involves using the Cartesian DDG application to generate three relaxed wild-type structures from the preprocessed experimental PDB structures for external datasets and AlphaFold2 structures for the Megascale dataset. The energies are subtracted from those from three replicate relaxed mutated structures, also generated by the application, in the REF2015 forcefield. We then use the three total score differences as three replicate ΔΔ*G* predictions without transformation or aggregation.

Predictions for ESM3 are generated almost identically to ESM-MSR. Unmasked wild-type and mutant sequences (accompanied by wild-type coordinates and structure tokens) are separately passed through the ESM3 models to obtain logits. Uncalibrated delta-logit scores from the two paths are averaged. Similarly, we generate ProteinMPNN predictions by encoding the wild-type structure alongside either the wild-type or mutant sequence. Because ProteinMPNN is an autoregressive model, we run M passes of the each sequence for an order-M multi-mutant, each with a different mutated position being decoded last such that the model has the full remaining sequence context. We sum the positional effects per sequence and then average over the two (wild-type and mutant) perspectives.

### External Datasets

#### Thermodynamic Folding Stability Datasets

We presented results from ten commonly used curated datasets of ΔΔ*G*; descriptions of these datasets can be found in their source material and in our previous work [37]: Q3421 [35], K2369 [18], Ssym [32], PTMUL and PTMUL(-D) [27], S461 [18], S2648 [7], S8754, S783 and S571 [48]. Although each dataset features mutations from several or potentially hundreds of proteins, characterization is very sparse and lacks the more comprehensive exploration of mutant effects provided by the Megascale dataset. Further, many datasets have overlapping entries (or are explicit subsets). Unlike Mutate Everything and ThermoMPNN-D, we used the full set of 912 mutations, de-duplicated from the 914 originally included for PTMUL. We used S461 instead of the more popular S669 superset, which is known to contain incorrect records and interface residues [18]. We use ΔΔ*G*_*u*_ with the sign convention of positive values representing higher mutant stability versus wild-type.

#### GRB2 & DLG4 Abundance & Binding

Faure et al. developed a “double deep protein complementation assay” (ddPCA) involving the functional complementation of fragments of dihydrofolate reductase (DHFR) [12]. Two protein domains were studied; in the authors’ own words: “the C-terminal SH3 domain of the human growth factor receptor-bound protein 2 (GRB2), which binds a proline-rich linear peptide of GRB2-associated binding protein 2 (GAB2), and the third PDZ domain from the adaptor protein PSD95/DLG4, which binds to the C-terminus of the protein CRIPT”. Both assays involve expressing fusion proteins with one of two complementary DHFR fragments fused to the studied domain. In the abundance assay, the other fragment is expressed at high levels in the cell, and restored DHFR activity via DHFR functional complementation indicates the level of expressed target domain present in the cell. The binding assay involves attaching the complementary fragment to the binding partner of each domain, such that DHFR activity indicates the frequency of binding between the two, and hence is related to both cellular abundance and binding affinity. The GRB2 dataset consists of 33,441 binding measurements and 63,366 abundance measurements, while the DLG4 dataset has 8,251 binding measurements and 6,976 abundance measurements. Both datasets contain a high fraction of possible single and double mutants. We used preprocessed datasets from ProteinGym but replaced the wild-type sequence with the truncated-structure sequence corresponding to the length of the studied domains. The values have been normalized such that 0 represents wild-type abundance and binding, and a larger value indicates higher abundance and binding.

#### Myoglobin Yeast Surface Display

Deep mutational scanning was performed on human myoglobin to search for stabilizing mutations, and Küng et al. showed that yeast surface display is an acceptable proxy for thermostability of human myoglobin [21]. We manually preprocessed the data using the notebooks provided by the authors, but deviated from their processing in that we did not filter out results where the codons began with A or C. Our preprocessed dataset contains 2,534 single mutations and 3,371 double mutants. A value of 0 represents wild-type level of display, and a larger value indicates greater display.

#### EstA ΔT_50_

Nutschel and colleagues studied the thermostability of single mutants of EstA lipase from *Bacillus subtilis* [30]. They measured *T*_50_, the temperature at which the enzyme showed 50% of its room temperature hydrolysis rate of para-nitrophenyl palmitate, estimated as the inflection point on a sigmoid fit of the activity versus temperature curve. We used the set of 2,170 single-mutants available from ProteinGym, which excludes many variants with stability effects smaller than the experimental error. A value of 0 represents wild-type *T*_50_, and a larger value indicates higher thermostability in terms of retained high-temperature activity, in Kelvin.

#### Human Domainome 1

Beltran and colleagues created a comprehensive database of cellular abundance for single mutants of human protein domains [2]. They used the same DHFR-based functional complementation assay as Faure et al. A value of 0 represents wild-type abundance and a larger value indicates higher abundance. We filtered the dataset to a total 536,145 unique mutations with valid structural positions from all 522 unique domains, reusing the structures from a previous work [22].

#### GB1 Binding Epistasis

Wu and colleages studied the binding of Streptococcal immunoglobulin-binding protein GB1 to the Fc region of IgG with nearly comprehensive combinatorial mutagenesis at 4 highly epistatic interacting sites (V39, D40, G41 and V54). The authors successfully characterized 149,360 out of a possible 159,999 single to quadruple mutants using their mRNA display pulldown assay methodology. Unlike the work of Faure et al., this binding assay measures binding affinity more directly, normalizing fitness using pre-pulldown read counts representing abundance in order to isolate enrichment due to binding [12].

### Data Analysis

#### Ranking Mutant Effects

We rely heavily on Spearman’s *ρ* to summarize mutant ranking performance. Aside from enabling robust, unbiased estimation of each model’s performance, it also enables ranking comparison without the assumption of a linear relationship, which is appropriately accommodates the likelihood-based models. Furthermore, this choice allows for meaningful analysis when the target property does not have units of kcal / mol, such as mutant abundance. Root-mean square deviation (RMSD) does not have these advantages, but we show our thermostability benchmark results when using this metric for completeness in the Supplementary Information, Figure 12.

#### Rectified Cumulative Gain at k

We introduced a new statistical measure designed to be maximally relevant and interpretable for the task of stability engineering with a known experimental budget. The rectified cumulative gain at k indicates the expected value of a screening experiment in physical units (kcal / mol). The value of selected destabilizing mutants is set to zero, while the value of stabilizing mutations is identical to their stabilizing effect (eq. (**??**). Rectifying the value is necessary to remove the confounding effects of very poor mutations from the total score. Highly destabilizing mutations are (to a first approximation) no less valuable than neutral ones in a traditional directed evolution campaign since only the most stabilizing mutants will be recombined in subsequent rounds.

This statistic does not account for the redundancy of multiple positive mutations at a single site.

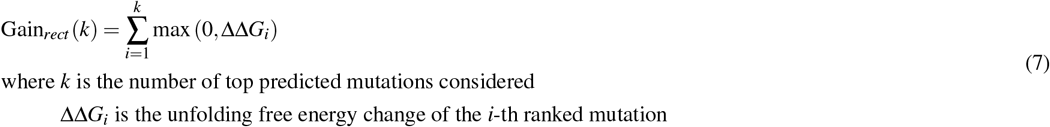

where *k* is the number of top predicted mutations considered ΔΔ*G*_*i*_ is the unfolding free energy change of the *i*-th ranked mutation

#### NDCG

We implemented normalized discounted cumulative gain (NDCG) in a very similar manner to rectified cumulative gain, setting the gain of destabilizing mutations to 0. While rectified gain is perhaps more interpretable for stability assays, NDCG is the traditional choice for protein engineering. NDCG at k establishes how well a model’s top k predictions capitalize on the property of interest, and is differentiated by discounting the importance of items lower in the ranked list. It is also normalized such that, in our case, 1 signifies perfectly ranked selection of the top k mutants, and 0 signifies selection of zero stabilizing mutants. In the Supplementary Information (Figure 13), we show our thermostability benchmark results evaluated using NDCG, but the conclusions largely match those when using Spearman’s *ρ*.

## Data Availability

All relevant data and code to reproduce figures and predictions are available on our GitHub page: https://github.com/SKTeamLab/esm-msr. Specifically, we provide pre-processing, inference, and analysis scripts to facilitate the reproduction of our analyses from the original source datasets made available by other authors. Raw predictions from all models analyzed are available on Zenodo for further scrutiny.

## Code Availability

LoRA weights for ESM-MSR as well as code for inference and reproducing the experiments and analyzes are available on GitHub under and MIT license: https://github.com/SKTeamLab/esm-msr. Our model depends on ESM-small-open-v1, whose weights were recently made publicly available under an MIT license and can be downloaded or accessed via HuggingFace: https://huggingface.co/biohub/esm3-sm-open-v1. Our repository also includes the ChimeraX tool that can be used to easily screen and visualize mutations for a loaded structure after some initial setup.

## Acknowledgments

The authors acknowledge the high-performance computing resources provided by the Digital Research Alliance of Canada. S.K. acknowledges the funding support from the NSERC Discovery Grant [RGPIN-2022-03348], NFRF Exploration Fund [NFRFE-2024-00791], and the Ontario Early Researcher Award [ER24-18-206]. S.R. acknowledges the NSERC CGS-D graduate scholarship.

## Author Contributions Statement

S.K. and S.R. conceived the project. S.R. designed and conducted the experiment(s), analyzed the results, and drafted the manuscript with input from S.K. S.K. supervised the project and acquired funding. All authors reviewed the manuscript.

