## Supplementary Information for "Robust Multi-Mutant Protein Stability Prediction from a Fine-Tuned Evolutionary Scale Model"

June 4, 2026

### *Ablations and Variations*

We tested the following ablations and variations of ESM-MSR and other models, shown in the subsequent 3 tables: ESM-MSR (as reported): Identical model(s) to main text, notably applies LoRA targeting the key, query and value projections (layernorm_qkv fused layers) and FFN up/down projection (ffn.1 & ffn.3) layers in all transformer blocks of the model. ESM-MSR, WT LoRA only.: uses only the wild-type sequence LoRA path, which does not encode mutant (non-additive) interactions. ESM-MSR, MT LoRA only.: uses only the mutant sequence LoRA path. ESM-MSR (indep. masking): One forward pass in computed through both LoRAs per mutated position, the mutated position is masked, and the per-position results are summed before averaging across paths as with the default model. ESM-MSR (masked marginal): One forward pass in computed through both LoRAs per mutant, with all mutated positions masked. ESM-MSR, ($\sigma = 1.25, 0.5, 0.25$): LoRA strength is multiplied by $\sigma$ for both LoRAs (default: $\alpha_{WT} = 4, \alpha_{MT} = 16$). ESM-MSR*, singles only: retrain exactly as originally, but exclude all double mutants. ESM-MSR*, no rank loss: retrained ablation where the ListMLE loss is not used, and the regression (MSE) loss is scaled up to keep the total loss similar. ESM-MSR*, detach reg. head: regression loss gradient detached from the LoRA, such that the LoRA is only responsible for learning relative ranks while the calibration head is only responsible for learning the scale and bias. ESM-MSR*, exclude seq. head: ablation where the final projection into sequence logits is not fine-tuned using LoRA. ESM-MSR*, target QKV, FFN, out_proj.: augmentation that additionally targets attention out projection layers for LoRA. ESM-MSR*, target QKV and out_proj.: variation that targets attention out projection layers and does not target FFN up/down projection layers. ESM-MSR*, target ffn and out_proj.: variation where LoRA is not applied to query, key and value projections , only attn_outproj and FFN layers. ESM-MSR*, only FFN up/down.: ablation where LoRA is only applied to FFN up/down projection layers. ESM-MSR*, only QKV proj.: ablation where LoRA is only applied to QKV projections. ESM-MSR*, WT&MT LoRA rank (1/4): LoRA rank is decreased from WT: 2, MT: 16 to either both 1 ($\alpha = 2$) or both 4 ($\alpha$=8). ESM-MSR*, WT&MT LoRA shared: the same LoRA ($r = 16, \alpha = 16$) is used for both the WT and MT paths. ESM-(small-open/small/medium/large): uses the same inference scheme as ESM-MSR (but with no LoRAs). ESM3-small-open (indep. masking / masked marginal): see analogous ESM-MSR versions above. ProteinMPNN: uses autoregressive decoding, where mutant positions are decoded last. Mutate Everything: trained according to the original method, on singles only for 20 epochs followed by singles and doubles for 100 epochs. Mutate Everything, singles only add. approx.: uses the intermediate singles only model from above and computes additive mutant effects using one forward pass per mutation, then summing scores. ThermoMPNN-D: trained according to the original method, up to 100 epochs on only double mutants with over-and-back augmentation with validation early stopping. ThermoMPNN, additive approx.: trained according to the original method, up to 100 epochs with validation early stopping on single mutants only and computes additive mutant effects using one forward pass per constituent mutation, then summing scores. SPURS-multi (retrained): trained by further fine-tuning the below model with a new script developed in consultation with the original authors including validation early stopping. No official training script was available upon request. SPURS (retrained) additive approx.: same as above, but trained with singles only and using the same additive approximation used by ThermoMPNN and MutateEverything. SPURS-multi, HuggingFace model: used the official SPURS-multi model out-of-the-box, trained on different splits from the remaining models. SPURS, HuggingFace model: used the official SPURS model out-of-the-box, trained on different splits from the remaining models and computes additive mutant effects.

**Table 1. Ranking Results for Ablations and Variations: Validation Set**. Spearman's $\rho$ is indicated for ungrouped (pooled) data and when scoring ranking of each domain's mutants before averaging over domains (grouped). Testing scaffolds include singles only (-S-), doubles only (-D-), combined, and $\delta\Delta\Delta G$ for doubles. All models are trained on the default split also used for hyperparameter optimization of ESM-MSR, except for the final two SPURS (HuggingFace) models (with double asterisk, **), zero-shot predictors, and Rosetta. Models with an asterisk (ESM-MSR*) are retrained with modifications, while those without use exactly the same model weights as ESM-MSR used in the main text. Highest performing models in each column are indicate in bold. ESM-MSR (as reported) is also bolded if the standard deviation range of its performance overlaps the highest average performance. Note that, for single mutations, the two masking strategies are equivalent.

| | hyperopt_splits-S-val | | hyperopt_splits-D-val | | hyperopt_splits-val | | hyperopt_splits-D-val-dddG | |
|---|---|---|---|---|---|---|---|---|
| ID | ungrouped | grouped | ungrouped | grouped | ungrouped | grouped | ungrouped | grouped |
| ESM-MSR (as reported) | **0.787 ± 0.003** | 0.823 ± 0.002 | **0.681 ± 0.017** | 0.702 ± 0.002 | **0.819 ± 0.003** | **0.836 ± 0.001** | 0.524 ± 0.009 | **0.497 ± 0.012** |
| ESM-MSR, WT LoRA only | 0.772 ± 0.004 | 0.800 ± 0.002 | 0.674 ± 0.019 | 0.614 ± 0.005 | 0.809 ± 0.002 | 0.808 ± 0.002 | 0.062 ± 0.067 | 0.012 ± 0.006 |
| ESM-MSR, MT LoRA only | 0.773 ± 0.003 | 0.817 ± 0.002 | 0.527 ± 0.016 | 0.667 ± 0.008 | 0.730 ± 0.005 | 0.797 ± 0.003 | 0.524 ± 0.009 | 0.497 ± 0.012 |
| ESM-MSR, indep. masking | 0.764 ± 0.004 | 0.810 ± 0.002 | 0.645 ± 0.018 | 0.689 ± 0.003 | 0.797 ± 0.004 | 0.822 ± 0.002 | 0.538 ± 0.021 | 0.501 ± 0.011 |
| ESM-MSR, masked marginal | 0.764 ± 0.004 | 0.810 ± 0.002 | 0.622 ± 0.013 | 0.645 ± 0.001 | 0.781 ± 0.004 | 0.814 ± 0.002 | 0.315 ± 0.019 | 0.311 ± 0.015 |
| ESM-MSR, $\sigma$=1.25 | **0.790 ± 0.002** | **0.826 ± 0.002** | 0.676 ± 0.020 | **0.712 ± 0.002** | 0.816 ± 0.004 | 0.838 ± 0.002 | 0.492 ± 0.006 | 0.499 ± 0.010 |
| ESM-MSR, $\sigma$=0.5 | 0.714 ± 0.003 | 0.755 ± 0.003 | 0.593 ± 0.002 | 0.602 ± 0.003 | 0.765 ± 0.002 | 0.779 ± 0.003 | 0.561 ± 0.009 | 0.466 ± 0.004 |
| ESM-MSR, $\sigma$=0.25 | 0.608 ± 0.002 | 0.663 ± 0.003 | 0.478 ± 0.001 | 0.468 ± 0.004 | 0.678 ± 0.001 | 0.695 ± 0.003 | 0.545 ± 0.004 | 0.411 ± 0.003 |
| ESM-MSR*, indep. masking | 0.770 ± 0.003 | 0.818 ± 0.001 | 0.646 ± 0.004 | 0.704 ± 0.006 | 0.800 ± 0.003 | 0.832 ± 0.001 | 0.573 ± 0.004 | 0.512 ± 0.015 |
| ESM-MSR*, masked marginal | 0.771 ± 0.005 | 0.814 ± 0.003 | 0.621 ± 0.014 | 0.691 ± 0.005 | 0.783 ± 0.003 | 0.822 ± 0.002 | 0.393 ± 0.039 | 0.347 ± 0.028 |
| ESM-MSR*, singles only | 0.788 ± 0.003 | 0.823 ± 0.002 | 0.670 ± 0.023 | 0.685 ± 0.006 | 0.818 ± 0.002 | 0.832 ± 0.001 | 0.462 ± 0.019 | 0.412 ± 0.017 |
| ESM-MSR*, no rank loss | 0.787 ± 0.003 | 0.812 ± 0.001 | 0.659 ± 0.007 | 0.691 ± 0.005 | 0.816 ± 0.001 | 0.825 ± 0.002 | 0.524 ± 0.018 | 0.475 ± 0.003 |
| ESM-MSR*, detach cal. head | 0.782 ± 0.001 | 0.822 ± 0.002 | 0.679 ± 0.003 | 0.703 ± 0.005 | 0.814 ± 0.002 | 0.835 ± 0.002 | 0.523 ± 0.007 | 0.492 ± 0.008 |
| ESM-MSR*, exclude seq. head | 0.788 ± 0.003 | 0.821 ± 0.001 | 0.689 ± 0.020 | 0.702 ± 0.006 | 0.821 ± 0.004 | 0.835 ± 0.001 | 0.518 ± 0.019 | 0.490 ± 0.006 |
| ESM-MSR*, target QKV, FFN, out_proj. | 0.786 ± 0.004 | 0.823 ± 0.001 | **0.691 ± 0.019** | 0.700 ± 0.004 | 0.819 ± 0.005 | **0.837 ± 0.001** | 0.526 ± 0.025 | **0.504 ± 0.013** |
| ESM-MSR*, target QKV and out_proj. | 0.779 ± 0.002 | 0.824 ± 0.001 | 0.652 ± 0.010 | 0.698 ± 0.004 | 0.802 ± 0.002 | 0.831 ± 0.001 | **0.539 ± 0.019** | 0.484 ± 0.009 |
| ESM-MSR*, target FFN and out_proj. | 0.777 ± 0.001 | 0.818 ± 0.002 | 0.665 ± 0.008 | 0.696 ± 0.001 | 0.809 ± 0.001 | 0.832 ± 0.001 | 0.518 ± 0.011 | 0.478 ± 0.001 |
| ESM-MSR*, only FFN up/down | 0.782 ± 0.005 | 0.819 ± 0.002 | 0.675 ± 0.020 | 0.701 ± 0.006 | 0.814 ± 0.007 | 0.833 ± 0.002 | 0.514 ± 0.010 | 0.486 ± 0.005 |
| ESM-MSR*, only QKV proj. | 0.771 ± 0.003 | 0.819 ± 0.001 | 0.636 ± 0.028 | 0.686 ± 0.002 | 0.796 ± 0.007 | 0.828 ± 0.001 | 0.489 ± 0.022 | 0.472 ± 0.016 |
| ESM-MSR*, WT&MT LoRA rank 1 | 0.770 ± 0.000 | 0.812 ± 0.001 | 0.659 ± 0.006 | 0.686 ± 0.002 | 0.802 ± 0.002 | 0.825 ± 0.001 | 0.466 ± 0.007 | 0.432 ± 0.005 |
| ESM-MSR*, WT&MT LoRA rank 4 | 0.782 ± 0.003 | 0.820 ± 0.000 | 0.661 ± 0.012 | 0.694 ± 0.003 | 0.812 ± 0.003 | 0.833 ± 0.001 | 0.524 ± 0.013 | 0.475 ± 0.011 |
| ESM-MSR*, WT+MT LoRA shared | 0.770 ± 0.000 | 0.812 ± 0.001 | 0.659 ± 0.006 | 0.686 ± 0.002 | 0.802 ± 0.002 | 0.825 ± 0.001 | 0.466 ± 0.007 | 0.432 ± 0.005 |
| ESM3-small-open | 0.487 | 0.524 | 0.388 | 0.265 | 0.579 | 0.561 | 0.475 | 0.319 |
| ESM3-small-open, indep. masking | 0.509 | 0.545 | 0.416 | 0.309 | 0.597 | 0.582 | 0.548 | 0.355 |
| ESM3-small-open, masked marginal | 0.509 | 0.545 | 0.401 | 0.279 | 0.590 | 0.576 | 0.300 | 0.229 |
| ESM3-small | 0.513 | 0.558 | 0.429 | 0.328 | 0.605 | 0.592 | 0.567 | 0.279 |
| ESM3-medium | 0.475 | 0.542 | 0.299 | 0.340 | 0.566 | 0.583 | 0.464 | 0.338 |
| ESM3-large | 0.431 | 0.476 | 0.373 | 0.255 | 0.556 | 0.516 | 0.442 | 0.243 |
| ProteinMPNN | 0.563 | 0.584 | 0.384 | 0.299 | 0.615 | 0.595 | 0.420 | 0.340 |
| Mutate Everything | 0.718 ± 0.005 | 0.729 ± 0.003 | 0.597 ± 0.026 | 0.576 ± 0.008 | 0.745 ± 0.003 | 0.743 ± 0.002 | 0.500 ± 0.052 | 0.246 ± 0.007 |
| Mutate Everything, singles only add. approx. | 0.732 ± 0.009 | 0.741 ± 0.008 | 0.662 ± 0.020 | 0.536 ± 0.007 | 0.774 ± 0.005 | 0.748 ± 0.005 | nan | nan |
| ThermoMPNN(-D) | 0.731 ± 0.002 | 0.757 ± 0.002 | 0.514 ± 0.009 | 0.569 ± 0.007 | 0.740 ± 0.010 | 0.762 ± 0.002 | 0.216 ± 0.021 | 0.161 ± 0.024 |
| ThermoMPNN, additive approx. | 0.731 ± 0.002 | 0.757 ± 0.002 | 0.619 ± 0.003 | 0.570 ± 0.007 | 0.766 ± 0.003 | 0.769 ± 0.001 | nan | nan |
| SPURS-multi (retrained) | 0.598 ± 0.014 | 0.600 ± 0.017 | 0.674 ± 0.012 | 0.577 ± 0.027 | 0.677 ± 0.024 | 0.640 ± 0.021 | 0.203 ± 0.072 | 0.241 ± 0.016 |
| SPURS (retrained), additive approx. | 0.687 ± 0.005 | 0.698 ± 0.006 | 0.609 ± 0.013 | 0.554 ± 0.003 | 0.738 ± 0.002 | 0.721 ± 0.004 | nan | nan |
| SPURS-multi, HuggingFace model** | 0.740 | 0.749 | 0.868 | 0.768 | 0.809 | 0.783 | 0.681 | 0.430 |
| SPURS, HuggingFace model add. approx.** | 0.851 | 0.865 | 0.758 | 0.641 | 0.861 | 0.843 | nan | nan |
| Rosetta Cartesian DDG | 0.628 ± 0.001 | 0.634 ± 0.001 | 0.620 ± 0.001 | 0.473 ± 0.001 | 0.676 ± 0.001 | 0.666 ± 0.001 | 0.367 ± 0.018 | 0.183 ± 0.015 |
| Rosetta, add. approx. | 0.628 ± 0.001 | 0.634 ± 0.001 | 0.537 ± 0.005 | 0.315 ± 0.003 | 0.669 ± 0.001 | 0.650 ± 0.000 | nan | nan |

**Table 2. Ranking Results for Ablations and Variations: Test Set**. Spearman's $\rho$ is indicated for ungrouped (pooled) data and when scoring ranking of each domain's mutants before averaging over domains (grouped). Testing scaffolds include singles only (-S-), doubles only (-D-), combined, and $\delta\Delta\Delta G$ for doubles. All models are trained on the default split also used for hyperparameter optimization of ESM-MSR, except for the final two SPURS (HuggingFace) models (with double asterisk, **), zero-shot predictors, and Rosetta. Models with an asterisk (ESM-MSR*) are retrained with modifications, while those without use exactly the same model weights as ESM-MSR used in the main text. Highest performing models in each column are indicate in bold. ESM-MSR (as reported) is also bolded if the standard deviation range of its performance overlaps the highest average performance. Note that, for single mutations, the two masking strategies are equivalent.

| | hyperopt_splits-S-test | | hyperopt_splits-D-test | | hyperopt_splits-test | | hyperopt_splits-D-test-dddG | |
|---|---|---|---|---|---|---|---|---|
| ID | ungrouped | grouped | ungrouped | grouped | ungrouped | grouped | ungrouped | grouped |
| ESM-MSR (as reported) | **0.774 ± 0.007** | **0.816 ± 0.002** | 0.357 ± 0.007 | 0.798 ± 0.003 | 0.727 ± 0.010 | **0.807 ± 0.003** | 0.263 ± 0.005 | **0.600 ± 0.005** |
| ESM-MSR, WT LoRA only | 0.770 ± 0.009 | 0.806 ± 0.005 | 0.339 ± 0.022 | 0.726 ± 0.006 | 0.762 ± 0.016 | 0.791 ± 0.005 | 0.029 ± 0.015 | 0.006 ± 0.062 |
| ESM-MSR, MT LoRA only | 0.752 ± 0.006 | 0.804 ± 0.002 | 0.319 ± 0.026 | 0.767 ± 0.010 | 0.597 ± 0.006 | 0.789 ± 0.002 | 0.263 ± 0.005 | 0.600 ± 0.005 |
| ESM-MSR, indep. masking | 0.766 ± 0.009 | 0.810 ± 0.003 | 0.278 ± 0.007 | 0.774 ± 0.006 | 0.669 ± 0.012 | 0.799 ± 0.003 | 0.264 ± 0.009 | 0.581 ± 0.013 |
| ESM-MSR, masked marginal | 0.766 ± 0.009 | 0.810 ± 0.003 | 0.348 ± 0.005 | 0.771 ± 0.005 | 0.662 ± 0.013 | 0.796 ± 0.003 | 0.189 ± 0.024 | 0.389 ± 0.015 |
| ESM-MSR, $\sigma$=1.25 | 0.763 ± 0.008 | 0.809 ± 0.003 | 0.343 ± 0.009 | 0.791 ± 0.006 | 0.703 ± 0.012 | 0.799 ± 0.004 | 0.230 ± 0.003 | 0.593 ± 0.010 |
| ESM-MSR, $\sigma$=0.5 | 0.758 ± 0.003 | 0.786 ± 0.004 | 0.343 ± 0.013 | 0.729 ± 0.011 | 0.740 ± 0.004 | 0.777 ± 0.004 | 0.276 ± 0.011 | 0.572 ± 0.003 |
| ESM-MSR, $\sigma$=0.25 | 0.693 ± 0.003 | 0.715 ± 0.004 | 0.242 ± 0.008 | 0.596 ± 0.009 | 0.686 ± 0.003 | 0.705 ± 0.004 | 0.265 ± 0.014 | 0.520 ± 0.004 |
| ESM-MSR*, indep. masking | 0.775 ± 0.002 | 0.814 ± 0.001 | 0.331 ± 0.037 | 0.781 ± 0.015 | 0.689 ± 0.011 | 0.802 ± 0.002 | 0.231 ± 0.014 | 0.559 ± 0.007 |
| ESM-MSR*, masked marginal | 0.769 ± 0.001 | 0.807 ± 0.002 | 0.379 ± 0.006 | 0.791 ± 0.006 | 0.677 ± 0.006 | 0.795 ± 0.001 | 0.172 ± 0.023 | 0.456 ± 0.014 |
| ESM-MSR*, singles only | 0.764 ± 0.001 | 0.812 ± 0.001 | 0.386 ± 0.006 | 0.803 ± 0.011 | 0.731 ± 0.007 | 0.802 ± 0.001 | 0.235 ± 0.008 | 0.515 ± 0.005 |
| ESM-MSR*, no rank loss | 0.773 ± 0.004 | 0.806 ± 0.003 | 0.371 ± 0.023 | 0.797 ± 0.005 | 0.717 ± 0.012 | 0.798 ± 0.004 | 0.269 ± 0.024 | 0.576 ± 0.005 |
| ESM-MSR*, detach cal. head | 0.767 ± 0.003 | 0.815 ± 0.002 | 0.378 ± 0.040 | 0.805 ± 0.006 | 0.719 ± 0.012 | 0.806 ± 0.003 | 0.260 ± 0.029 | 0.589 ± 0.007 |
| ESM-MSR*, exclude seq. head | 0.768 ± 0.002 | 0.813 ± 0.002 | 0.371 ± 0.045 | 0.803 ± 0.006 | 0.724 ± 0.018 | 0.804 ± 0.003 | 0.272 ± 0.009 | 0.596 ± 0.008 |
| ESM-MSR*, target QKV, FFN, out_proj. | **0.775 ± 0.004** | 0.815 ± 0.003 | 0.403 ± 0.034 | **0.806 ± 0.007** | 0.733 ± 0.007 | 0.807 ± 0.003 | 0.258 ± 0.025 | 0.581 ± 0.012 |
| ESM-MSR*, target QKV and out_proj. | 0.772 ± 0.005 | 0.813 ± 0.002 | 0.382 ± 0.008 | 0.802 ± 0.007 | 0.728 ± 0.007 | 0.806 ± 0.003 | 0.239 ± 0.031 | 0.590 ± 0.014 |
| ESM-MSR*, target FFN and out_proj. | **0.775 ± 0.001** | 0.813 ± 0.001 | 0.405 ± 0.030 | 0.798 ± 0.006 | 0.735 ± 0.007 | 0.804 ± 0.001 | 0.242 ± 0.025 | 0.575 ± 0.021 |
| ESM-MSR*, only FFN up/down | 0.769 ± 0.002 | 0.813 ± 0.001 | 0.355 ± 0.038 | 0.801 ± 0.005 | 0.724 ± 0.007 | 0.803 ± 0.001 | 0.244 ± 0.009 | 0.587 ± 0.007 |
| ESM-MSR*, only QKV proj. | 0.768 ± 0.005 | 0.814 ± 0.001 | 0.348 ± 0.025 | 0.794 ± 0.008 | 0.712 ± 0.016 | **0.808 ± 0.002** | 0.249 ± 0.029 | **0.604 ± 0.005** |
| ESM-MSR*, WT&MT LoRA rank 1 | 0.763 ± 0.005 | 0.805 ± 0.005 | 0.369 ± 0.019 | 0.795 ± 0.003 | 0.706 ± 0.003 | 0.796 ± 0.006 | 0.268 ± 0.019 | 0.579 ± 0.006 |
| ESM-MSR*, WT&MT LoRA rank 4 | 0.773 ± 0.003 | 0.815 ± 0.001 | 0.408 ± 0.017 | 0.810 ± 0.008 | 0.738 ± 0.006 | 0.807 ± 0.002 | 0.263 ± 0.009 | 0.584 ± 0.003 |
| ESM-MSR*, WT+MT LoRA shared | 0.763 ± 0.005 | 0.805 ± 0.005 | 0.369 ± 0.019 | 0.795 ± 0.003 | 0.706 ± 0.003 | 0.796 ± 0.006 | 0.268 ± 0.019 | 0.579 ± 0.006 |
| ESM3-small-open | 0.546 | 0.561 | 0.105 | 0.419 | 0.546 | 0.551 | 0.232 | 0.413 |
| ESM-small-open, indep. masking | 0.559 | 0.579 | 0.089 | 0.422 | 0.536 | 0.570 | 0.207 | 0.419 |
| ESM-small-open, masked marginal | 0.559 | 0.579 | 0.108 | 0.427 | 0.530 | 0.570 | 0.242 | 0.314 |
| ESM3-small | 0.598 | 0.614 | 0.133 | 0.432 | 0.591 | 0.606 | 0.391 | 0.439 |
| ESM3-medium | 0.598 | 0.600 | 0.425 | 0.378 | 0.709 | 0.590 | 0.471 | 0.446 |
| ESM3-large | 0.539 | 0.555 | 0.460 | 0.402 | 0.681 | 0.553 | 0.351 | 0.367 |
| ProteinMPNN | 0.546 | 0.539 | 0.211 | 0.426 | 0.597 | 0.529 | 0.251 | 0.436 |
| Mutate Everything | 0.686 ± 0.009 | 0.693 ± 0.009 | 0.523 ± 0.051 | 0.637 ± 0.010 | 0.713 ± 0.018 | 0.685 ± 0.011 | **0.467 ± 0.056** | 0.352 ± 0.066 |
| Mutate Everything, singles only additive approx. | 0.696 ± 0.015 | 0.703 ± 0.017 | 0.577 ± 0.032 | 0.633 ± 0.030 | **0.786 ± 0.012** | 0.700 ± 0.017 | nan | nan |
| ThermoMPNN(-D) | 0.729 ± 0.003 | 0.745 ± 0.003 | 0.505 ± 0.026 | 0.730 ± 0.012 | 0.716 ± 0.008 | 0.735 ± 0.005 | 0.176 ± 0.100 | 0.235 ± 0.035 |
| ThermoMPNN, additive approx. | 0.729 ± 0.003 | 0.745 ± 0.003 | 0.507 ± 0.007 | 0.724 ± 0.011 | 0.769 ± 0.005 | 0.737 ± 0.003 | nan | nan |
| SPURS-multi (retrained) | 0.606 ± 0.013 | 0.626 ± 0.017 | 0.591 ± 0.055 | 0.670 ± 0.051 | 0.714 ± 0.004 | 0.633 ± 0.020 | 0.179 ± 0.135 | 0.236 ± 0.100 |
| SPURS (retrained), additive approx. | 0.680 ± 0.014 | 0.703 ± 0.013 | **0.593 ± 0.008** | 0.695 ± 0.019 | 0.763 ± 0.007 | 0.704 ± 0.014 | nan | nan |
| SPURS-multi, HuggingFace model** | 0.720 | 0.732 | 0.586 | 0.769 | 0.779 | 0.735 | 0.561 | 0.507 |
| SPURS, HuggingFace model add. approx**. | 0.797 | 0.805 | 0.571 | 0.713 | 0.843 | 0.792 | nan | nan |
| Rosetta Cartesian DDG | 0.621 ± 0.001 | 0.645 ± 0.001 | 0.482 ± 0.001 | 0.467 ± 0.005 | 0.657 ± 0.000 | 0.634 ± 0.001 | 0.321 ± 0.012 | 0.225 ± 0.014 |
| Rosetta, add. approx. | 0.621 ± 0.001 | 0.645 ± 0.001 | 0.391 ± 0.002 | 0.362 ± 0.012 | 0.646 ± 0.000 | 0.615 ± 0.001 | nan | nan |

**Table 3. Ranking Results for Ablations and Variations: Domainome**. Spearman's $\rho$ is indicated for ungrouped (pooled) data and when scoring ranking of each domain's mutants before averaging over domains (grouped). All models are trained on the default split also used for hyperparameter optimization of ESM-MSR, except for the final two SPURS (HuggingFace) models (with double asterisk, **), zero-shot predictors, and Rosetta. Models with an asterisk (ESM-MSR*) are retrained with modifications, while those without use exactly the same model weights as ESM-MSR used in the main text. Highest performing models in each column are indicate in bold. ESM-MSR (as reported) is also bolded if the standard deviation range of its performance overlaps the highest average performance. Rosetta is not included due to compute constraints. Note that, for single mutations, the two masking strategies are equivalent.

| | domainome | |
|---|---|---|
| ID | ungrouped | grouped |
| ESM-MSR (as reported) | 0.546 ± 0.002 | 0.549 ± 0.003 |
| ESM-MSR, WT LoRA only | 0.545 ± 0.003 | 0.548 ± 0.004 |
| ESM-MSR, MT LoRA only | 0.524 ± 0.001 | 0.530 ± 0.001 |
| ESM-MSR, indep. masking | 0.556 ± 0.002 | 0.565 ± 0.003 |
| ESM-MSR, masked marginal | 0.556 ± 0.002 | 0.565 ± 0.003 |
| ESM-MSR, $\sigma$=1.25 | 0.525 ± 0.002 | 0.527 ± 0.004 |
| ESM-MSR, $\sigma$=0.5 | **0.564 ± 0.001** | **0.573 ± 0.001** |
| ESM-MSR, $\sigma$=0.25 | 0.550 ± 0.001 | 0.568 ± 0.001 |
| ESM-MSR*, indep. masking | 0.541 ± 0.004 | 0.550 ± 0.005 |
| ESM-MSR*, masked marginal | 0.534 ± 0.007 | 0.539 ± 0.007 |
| ESM-MSR*, singles only | 0.548 ± 0.003 | 0.549 ± 0.002 |
| ESM-MSR*, no rank loss | 0.536 ± 0.005 | 0.540 ± 0.007 |
| ESM-MSR*, detach cal. head | 0.541 ± 0.008 | 0.544 ± 0.009 |
| ESM-MSR*, exclude seq. head | 0.544 ± 0.002 | 0.547 ± 0.002 |
| ESM-MSR*, target QKV, FFN, out_proj. | 0.538 ± 0.003 | 0.542 ± 0.004 |
| ESM-MSR*, target QKV and out_proj. | 0.534 ± 0.005 | 0.539 ± 0.006 |
| ESM-MSR*, target FFN and out_proj. | 0.536 ± 0.004 | 0.539 ± 0.005 |
| ESM-MSR*, only FFN up/down | 0.541 ± 0.004 | 0.543 ± 0.004 |
| ESM-MSR*, only QKV proj. | 0.534 ± 0.002 | 0.540 ± 0.004 |
| ESM-MSR*, WT&MT LoRA rank 1 | 0.528 ± 0.005 | 0.534 ± 0.006 |
| ESM-MSR*, WT&MT LoRA rank 4 | 0.542 ± 0.004 | 0.546 ± 0.003 |
| ESM-MSR*, WT+MT LoRA shared | 0.528 ± 0.005 | 0.534 ± 0.006 |
| ESM3-small-open | 0.508 | 0.533 |
| ESM-small-open, indep. masking | 0.527 | 0.550 |
| ESM-small-open, masked marginal | 0.527 | 0.550 |
| ESM3-small | 0.529 | 0.547 |
| ESM3-medium | 0.503 | 0.531 |
| ESM3-large | 0.456 | 0.508 |
| ProteinMPNN | 0.531 | 0.547 |
| Mutate Everything | 0.523 ± 0.001 | 0.517 ± 0.001 |
| ThermoMPNN | 0.467 ± 0.005 | 0.455 ± 0.007 |
| SPURS-multi (retrained) | 0.303 ± 0.029 | 0.279 ± 0.035 |
| SPURS (retrained), additive approx. | 0.402 ± 0.005 | 0.376 ± 0.009 |
| SPURS-multi, HuggingFace model** | 0.412 | 0.406 |
| SPURS, HuggingFace model add. approx.** | 0.515 | 0.513 |

### *Computation Complexity of Screening*

**Table 4. Computational Complexity of Mutant Inference Strategies**. For full mutational scans, $N = L$ (sequence length). $N$: Positions to scan; $D$: Mutation depth e.g. 2 for a double mutant screen, $K$: Mutations to assess per position (e.g. 19); $T_{fwd}$: Forward pass time. The screening time assumes a protein of length 200, a batch size of 16, and a (conservatively high) estimated forward pass time of 1 second. Note that a forward pass may require more time for larger proteins. All results in the main text use the Unmasked strategy.

| Strategy | Single Mutants / Additive approx. (WT LoRA) | Time to Screen All Singles (L=200) | Multi-mutants / Epistatic approach (Dual-view) | Double Mutant Forward Passes (L=200) | Time to Screen All Doubles (L=200) | Notes |
|---|---|---|---|---|---|---|
| **Unmasked (Default)** | $1 \times T_{fwd}$ | 1 Second | $\left(\binom{N}{D} \cdot K^D + 1\right) \cdot T_{fwd}$ | 7,183,901 | 5.2 Days | Can be reduced by an order of magnitude by restricting inference to nearby residues. |
| **Independent Masking** | $N \times T_{fwd}$ | 12.5 Seconds | $\left(N + \binom{N}{D-1} \cdot K^{D-1}(N-D+1)\right) \cdot T_{fwd})$ | 756,400 | 13.1 Hours | Minimal performance loss versus unmasked. |
| **Masked Marginal** | $N \times T_{fwd}$ | 12.5 Seconds | $\binom{N}{D} \cdot T_{fwd}$ | 19,900 | 20.7 Minutes | Does not account for identity-dependent epistasis. |

**Table 5.** Total Compute Time in Seconds for Benchmark Categories

| | Megascale | Thermostability | Functional DMS | Domainome |
|---|---|---|---|---|
| ESM-MSR | 643 | 1928 | 1693 | 3666 |
| ESM-MSR (WT only) | 43 | 206 | 121 | 282 |
| ESM-MSR (chain) | 215 | 2073 | 1492 | 1197 |
| ESM-MSR (marginal) | 211 | 1943 | 842 | 1257 |

***Training and Testing Scaffold Information***

**Table 6. Summary of Training and Test Data**. Note that the test set contains two destabilized backbone variants of PDB ID 2K5H, which are counted in the main as different domains, hence 36 unique base sequences with 34 unique structures. For Ssym, each mutation has a unique structure, reducing the apparent mutations per protein.

| Superset | Unique Structures | Unique Mutations | Mutations per Protein |
|---|---|---|---|
| Megascale - Train | 120 | 183379 | 1528.16 |
| Megascale - Validation | 26 | 35373 | 1360.50 |
| Megascale - Test | 34 | 43700 | 1285.29 |
| Ssym | 357 | 684 | 1.92* |
| PTMUL | 90 | 912 | 10.13 |
| S461 | 48 | 461 | 9.60 |
| K2369 | 181 | 2369 | 13.09 |
| Q3421 | 149 | 3421 | 22.96 |
| S571 | 39 | 557 | 14.28 |
| S783 | 55 | 783 | 14.24 |
| S2648 | 132 | 2648 | 20.06 |
| S8754 | 305 | 6400 | 20.98 |
| DLG4 Abundance | 1 | 6976 | 6976 |
| DLG4 Binding | 1 | 8251 | 8251 |
| GRB2 Abundance | 1 | 63366 | 63366 |
| GRB2 Binding | 1 | 33441 | 33441 |
| MYO Display | 1 | 5905 | 5905 |
| ESTA dTm | 1 | 2172 | 2172 |
| GB1 Binding | 5 | 149360 | 149360 |
| Domainome | 522 | 536145 | 1027.10 |

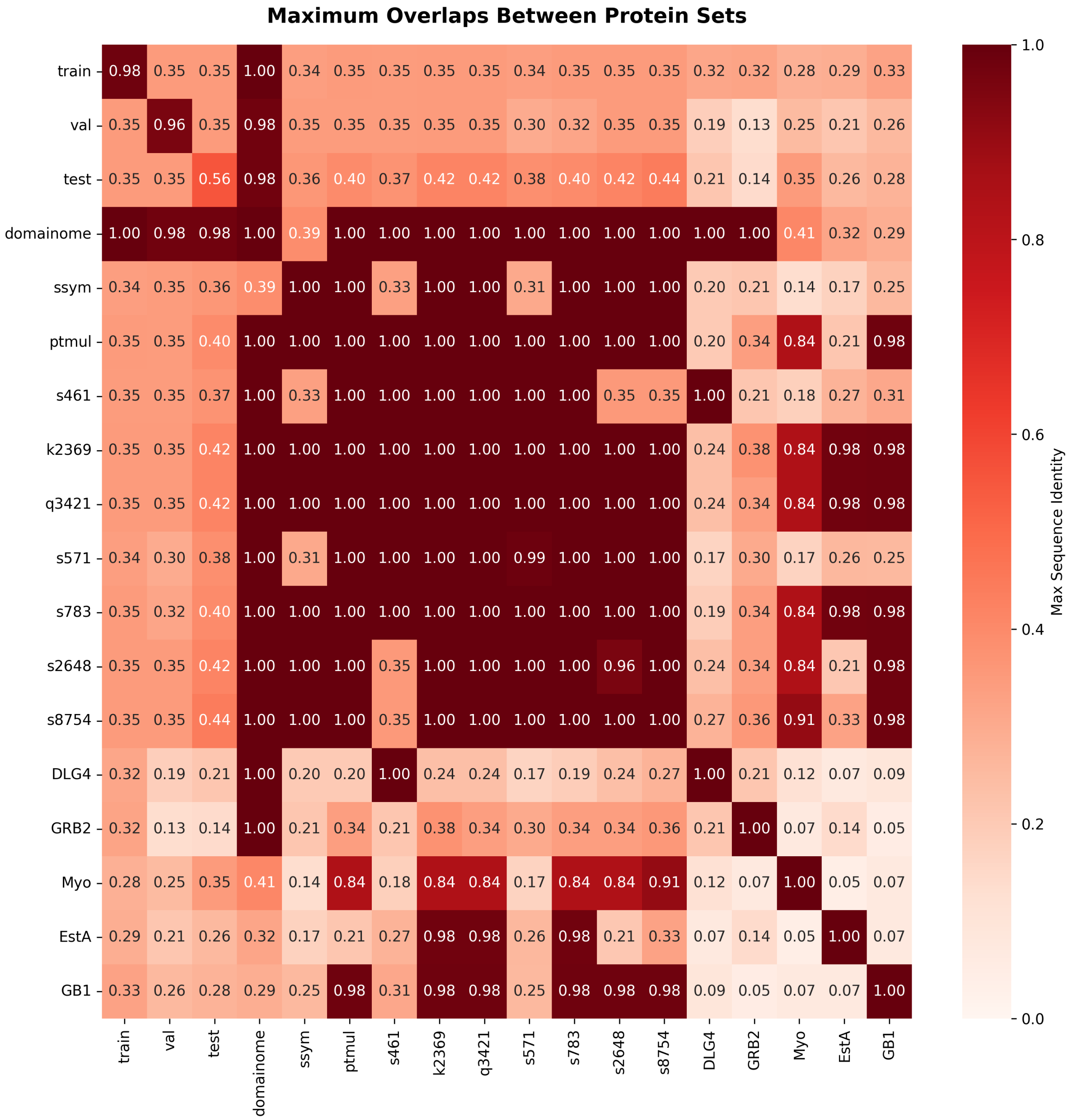

**Figure 1. Sequence Identity Overlaps between Training, Validation and Test Sets**. Maximum pairwise aligned percent identity between sets, defined for a given pair as number of matches divided by minimum sequence length.

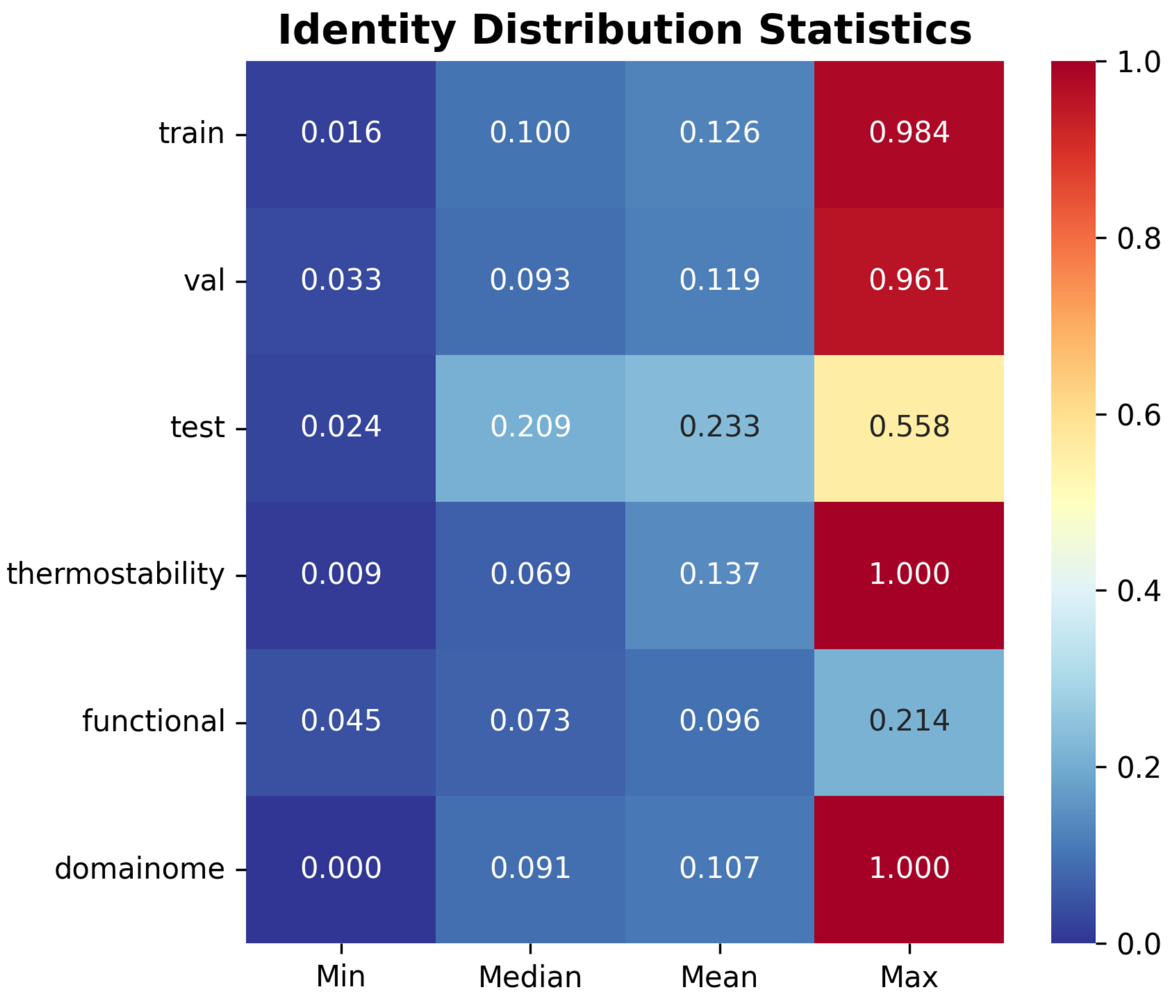

**Figure 2.** **Sequence Identity Overlaps within Each Set**.

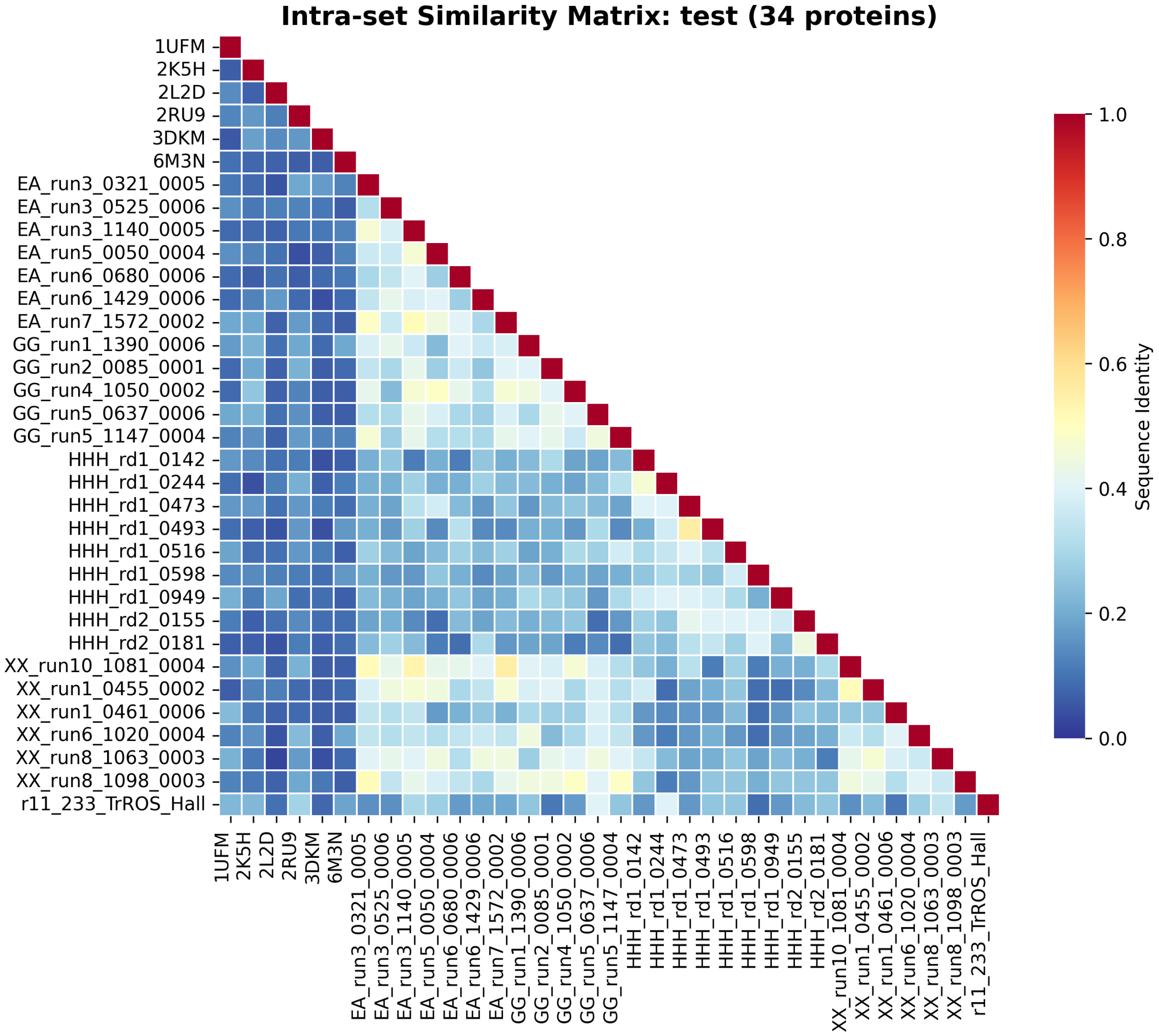

**Figure 3.** **Self-Similarity of the Test Set**.

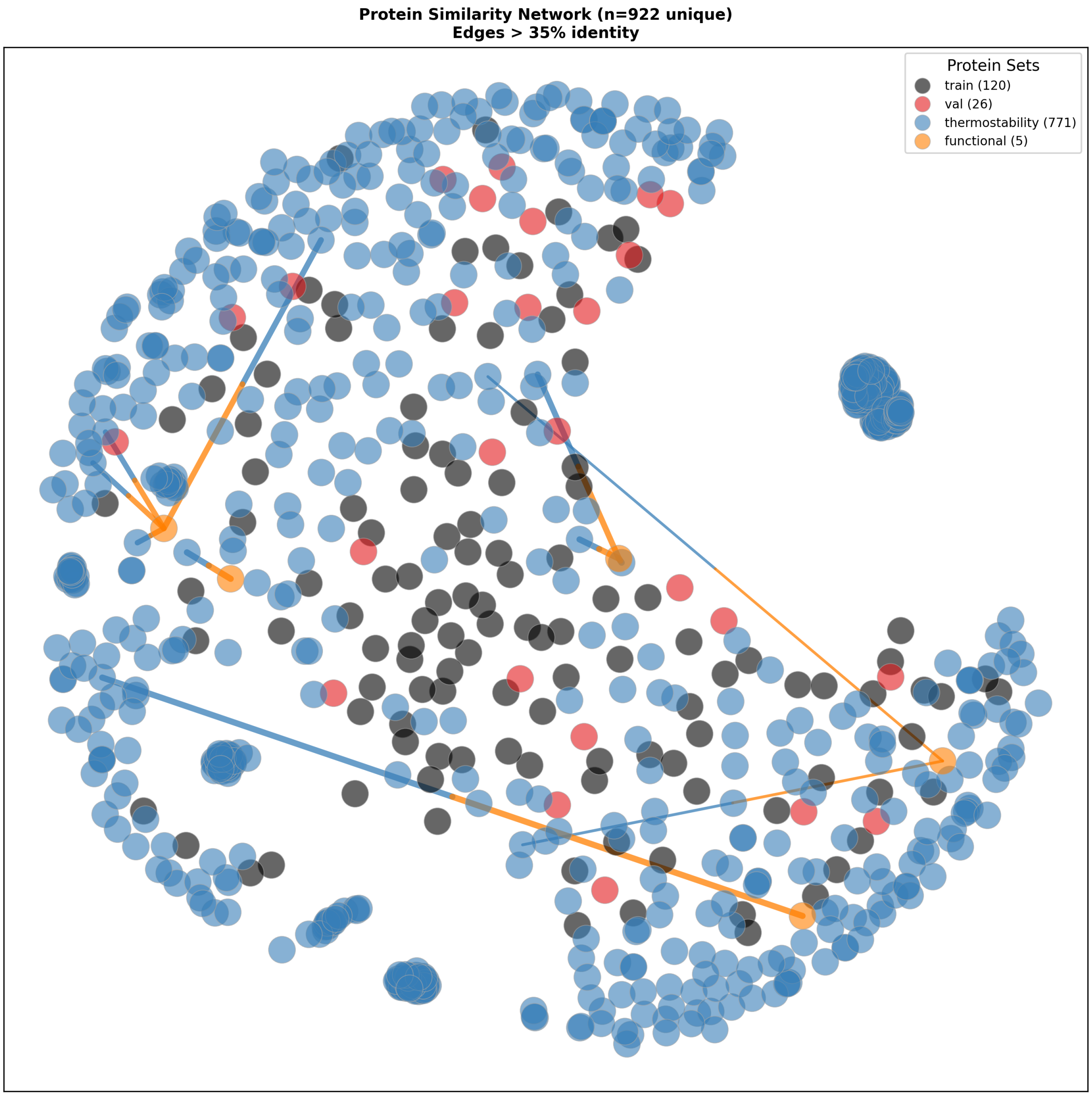

**Figure 4. Sequence Identity Overlaps between Training and External Datasets**. Dashed lines indicate intra-set identities exceeding 35%, while solid (two-colored) lines indicate inter-set overlap (filter violations). Visualized distances are based on the multidimensional scaling (MDS) algorithm. Even if two circles appear to overlap, they only exceed 35% identity (and are from different scaffolds) if they are connected by a line. Dense clusters of blue correspond to many structures one mutation from the wild-type in Ssym.

### *Assay Floor Limitations*

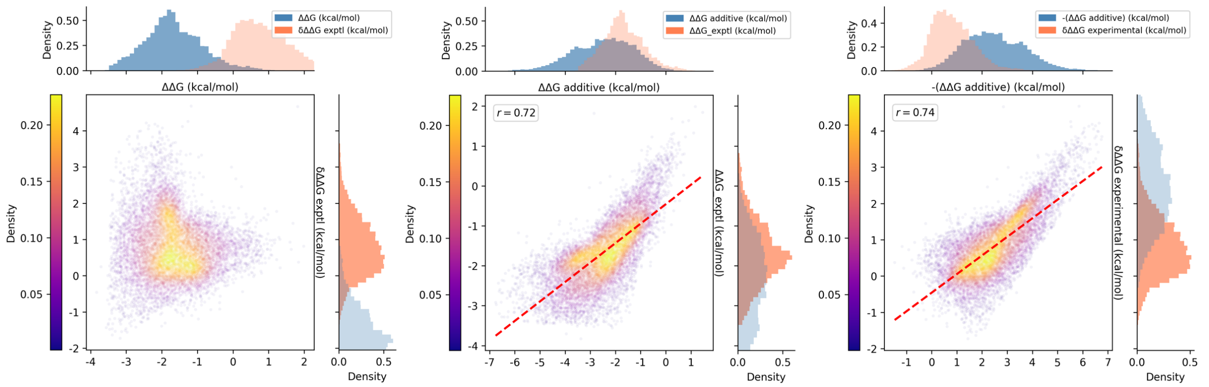

**Figure 5. Comparison of Distributions for Stability and Epistasis**. Data shown is for double mutants in the test set. The assay floor appears at approximately -3.5 kcal/mol for $\Delta\Delta G$. Experimental epistasis is mainly positive and rarely below 1 kcal/mol.

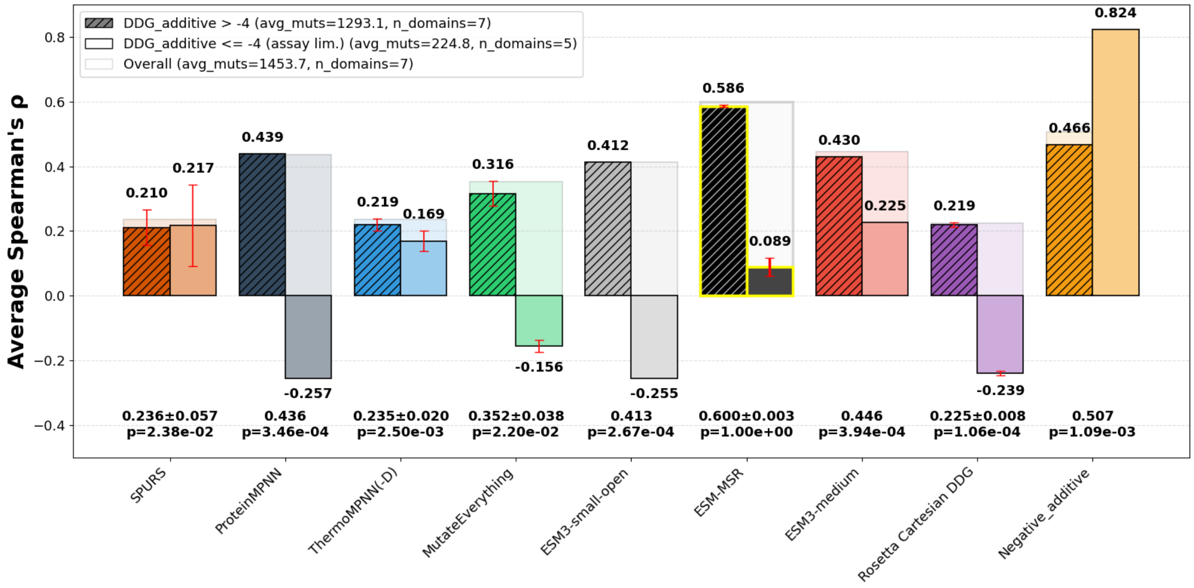

**Figure 6. Epistasis Prediction Dependence on Assay Limitations**. Shown above, the lowest measurable $\Delta\Delta G$ is approximately -3.5 kcal/mol. If mutations have additive effects below this threshold, they necessarily exhibit compensatory epistasis. As shown in Main Figure 2a (and here), this causes the negative correlation between $\delta\Delta\Delta G$ and additive $\Delta\Delta G$ to strengthen significantly below this point, leading to huge variance in model performance.

### *Canonical Megascale Split Results*

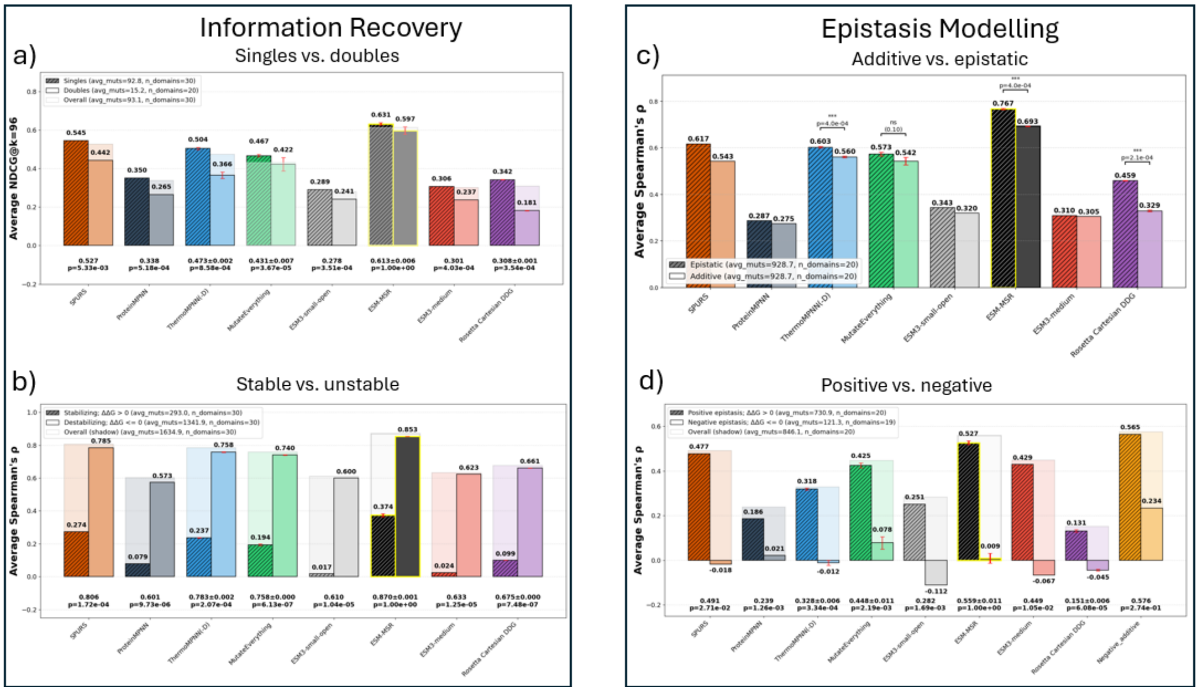

**Figure 7.** **Comparison of Models Using Canonical Megascale Split.** This figure seeks to fairly assess SPURS by using the official model and retraining and testing the others on the original Megascale splits from Dieckhaus et al. a): Test set information recovery performance for the full set of 49,048 test set mutations (including those with destabilized backbones, which were not originally assessed), overall (background shadow / foreground ghost) and after splitting into single (hatched bars) and double (solid bars) mutant subsets. Average $NDCG@96$ uses the top 96 highest scoring mutants per model and assigns gains for stabilizing mutants only (otherwise, 0). Some domains have fewer than 96 true stabilizing mutants, tracked in avg_muts in the legend. For transferred models, error bars represent the standard deviation across three differently seeded training runs using the same data. For Rosetta, they represent the standard deviation in score across three pairs of mutant and wild-type structure models. $p$ values represent the probability of the null hypothesis that the overall performance is the same as ESM-MSR; models with error bars are compared to ESM-MSR using a two-sample Welch's t-test, while pre-trained models have only one sample and use a one-sided t-test. b)-d): Test set ranking performance (average Spearman's $\rho$), using analogous visual elements. b) All test mutations, overall or split into stabilizing ($\Delta\Delta G > 0$) or destabilizing ($\Delta\Delta G \leq 0$) subsets. c) Double mutants only, split into epistatic (native double mutant predictions) or additive approximation (sum of constituent mutation predictions). d) For 16,922 double mutants with both singles characterized, epistasis is derived from experimental or predicted $\Delta\Delta G_{AB} - \Delta\Delta G_A - \Delta\Delta G_B$, overall and split into positive ($\delta\Delta\Delta G > 0$) or negative ($\delta\Delta\Delta G \leq 0$) epistasis subsets.

*Epistasis Classification*

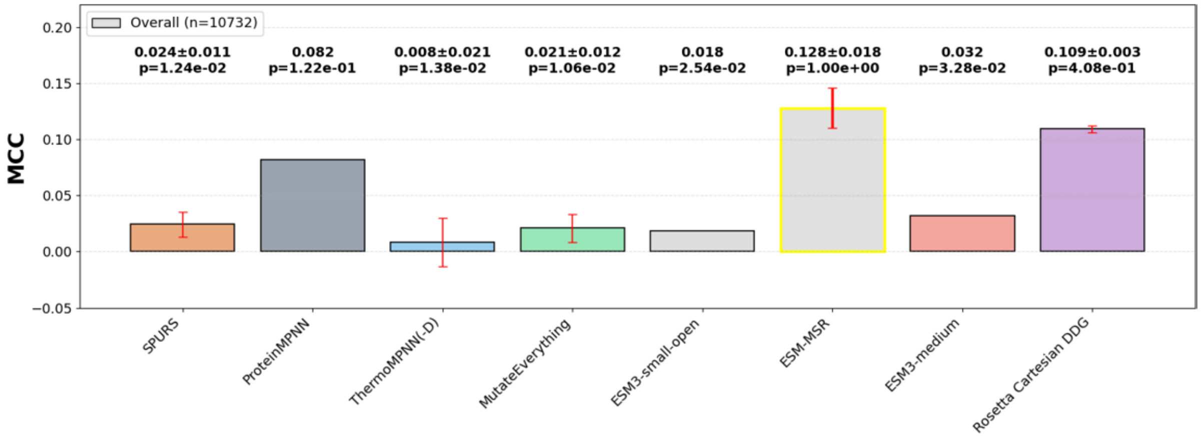

**Figure 8. Classification of Positive Versus Negative Epistasis**. Matthew's correlation coefficient for distinguish positive versus negative epistasis at a predicted and ground truth class threshold of 0 kcal/mol. Error bars represent the standard deviation across three training runs with different seeds, except for Rosetta, where it instead uses three stochastically generated mutant structures.

***Importance of the Domain Type***

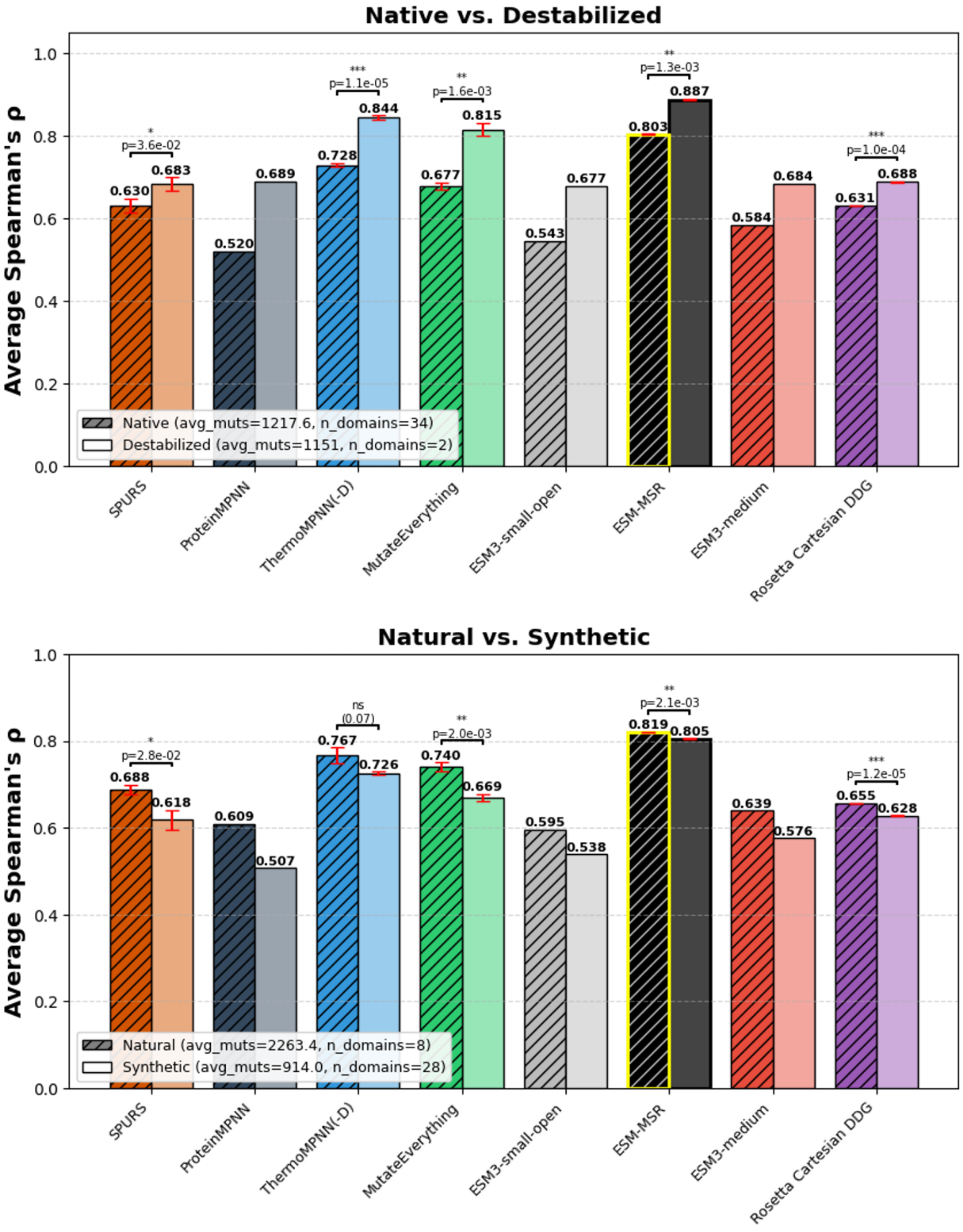

**Figure 9.** **Comparison of Models on Different Mutation Types.** In each subplot, mutations in the test set are split into two categories. We reuse the main training splits with 3 replicates per trained model; error bars represent the standard deviation in average Spearman's $\rho$ across replicates. Top: The scores are computed for all mutants derived from native and destabilized domains from the main test set, separately. Bottom: The scores are computed for all mutants derived from natural and synthetic (*de novo* designed) domains.

### ***Mutational Preferences for k=24***

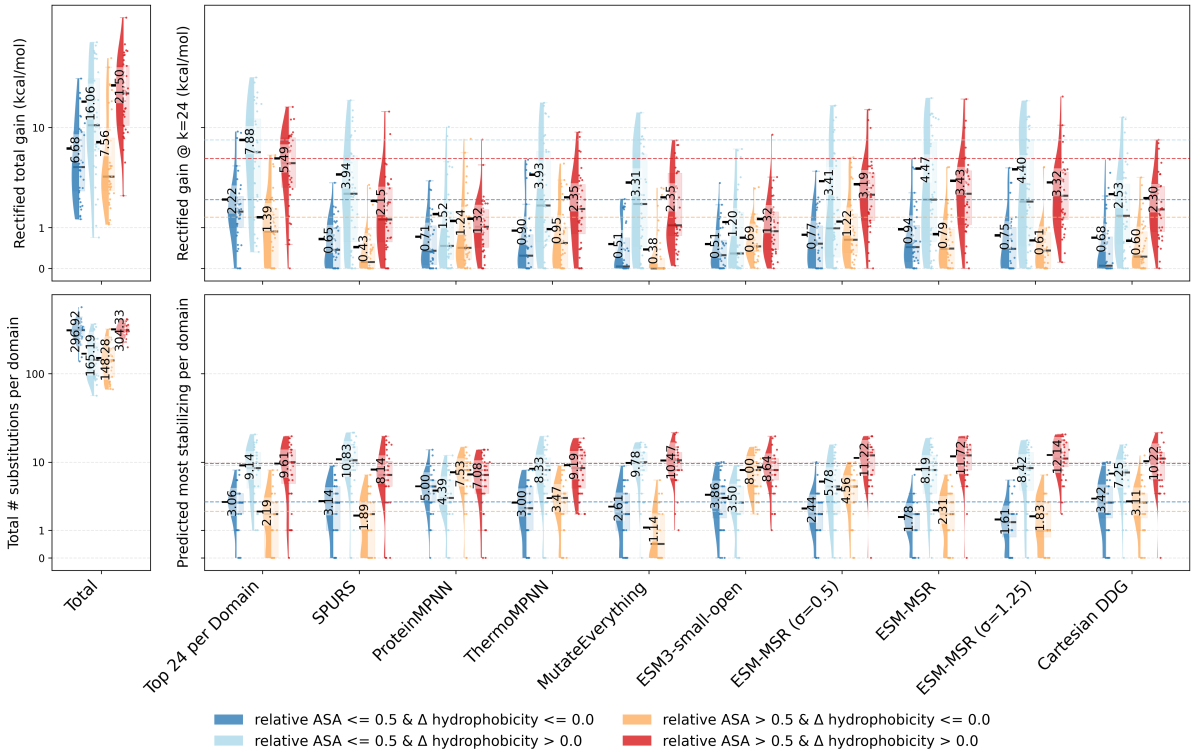

**Figure 10.** **Physicochemical Characteristics of Top 24 High-Scoring Mutations**. Identical to Figure 3b in the main text, but with 24 selected mutations instead of 96.

### *Ssym Antisymmetry and Bias*

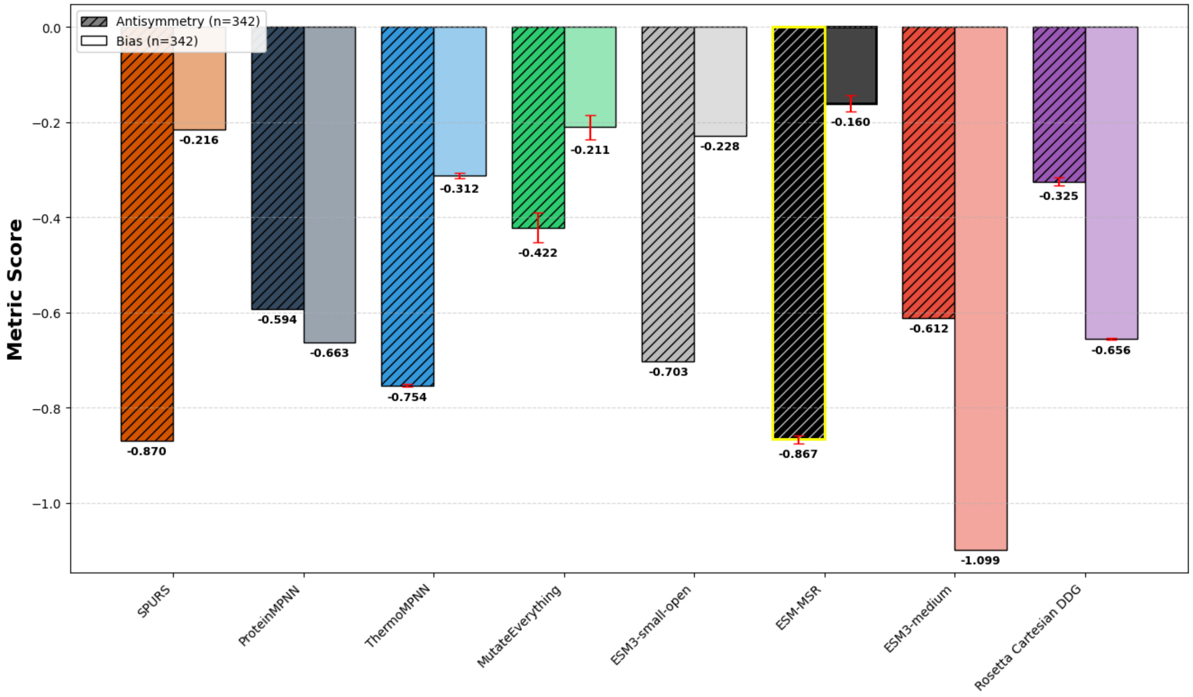

**Figure 11. Antisymmetry and Bias on the Ssym Dataset.** Forward and reverse predictions for 342 matched substitutions and reversions from Ssym are compared. Antisymmetry is optimized at -1 (forward predictions are of the same magnitude and opposite sign as reverse), while bias is optimized at 0 (the whole set of predictions is centered at 0).

### *Alternative Test Statistic: Root Mean Squared Error*

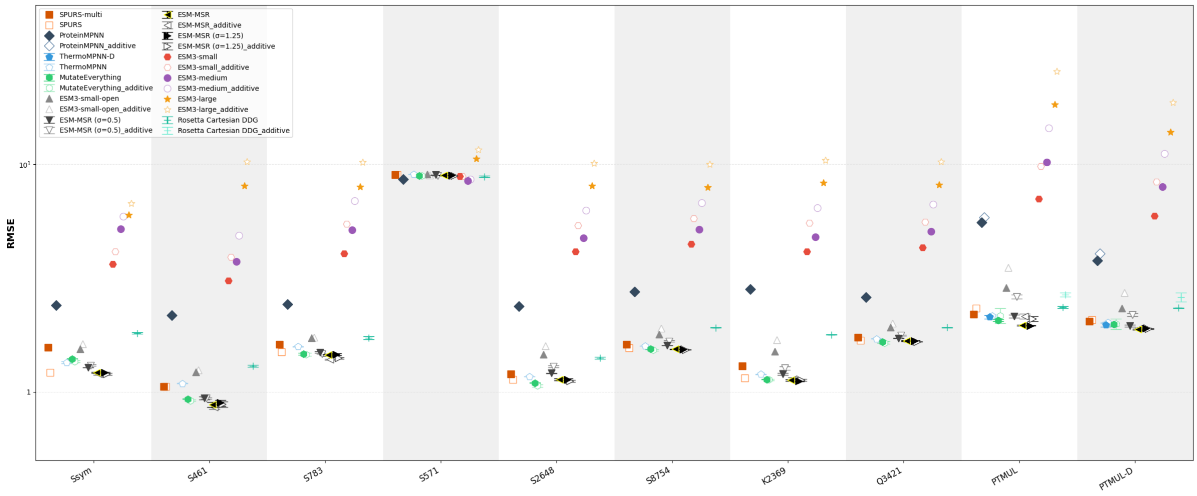

**Figure 12. Prediction Error of Stability Predictors on External Datasets**. Identical to Figure 5a in the main text, but the selected statistic is changed to the root mean squared error (RMSE), plotted on a log scale. Error bars represent one standard deviation around the mean computed from 3 replicate models trained on the same scaffold of the Megascale dataset. White markers with colored outlines represent the additive approximations corresponding to original models sharing the marker shape. For single mutants, these predictions are the same. For ThermoMPNN-D, ThermoMPNN predictions are used for single- and multi-mutants, since ThermoMPNN-D can only predict double mutants.

### *Alternative Test Statistic: NDCG*

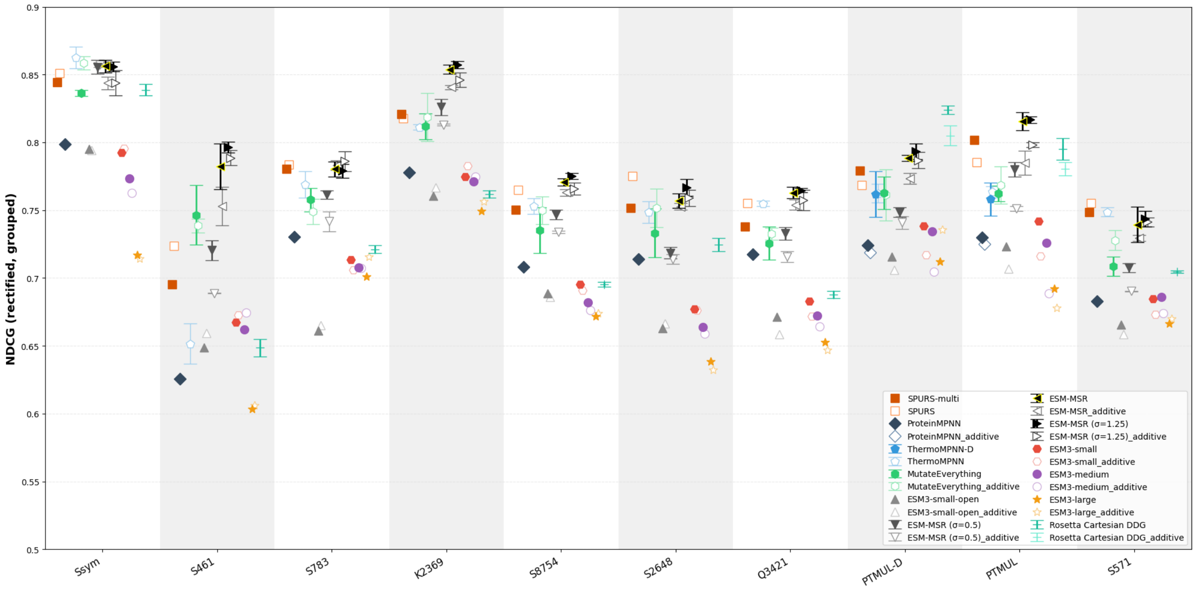

**Figure 13. Information Retrieval Performance of Stability Predictors on External Datasets**. Identical to Figure 5b from the main text, but the selected statistic is changed to the NDCG of all stabilizing mutations.
